# Thiol methyltransferase METTL7B coordinates lipid metabolism and promotes tumor progression in pancreatic cancer

**DOI:** 10.64898/2026.04.09.717095

**Authors:** Shuhei Mitsutomi, Daisuke Saigusa, Nobue Tamamura, Hiroki Tanaka, Anzu Sugawara, Atsuko Nakanishi-Ozeki, Kenji Takahashi, Yusuke Mizukami, Junken Aoki, Kenzui Taniue, Nobuyoshi Akimitsu

## Abstract

Dysregulated lipid metabolism drives cancer progression, yet the molecular mechanisms linking lipid accumulation to pancreatic cancer remain poorly understood. Here, we identify methyltransferase-like 7B (METTL7B) as a key regulator of lipid metabolism and tumor progression in pancreatic cancer. Integrated analysis of large-scale transcriptomic datasets revealed that METTL7B is upregulated in pancreatic cancer compared with normal tissues, and elevated METTL7B expression is associated with poor patient prognosis. Functional experiments demonstrated that METTL7B is required for pancreatic cancer cell proliferation, migration, and metastasis. We identified hepatocyte nuclear factor 4α (HNF4A) and hepatocyte nuclear factor 4γ (HNF4G), well-established regulators of lipid homeostasis, as transcription factors that directly upregulate METTL7B expression in pancreatic cancer cells. Moreover, we showed that METTL7B localizes around lipid droplets and promotes their accumulation in pancreatic cancer cells. Lipidomic analyses revealed that METTL7B depletion selectively reduces long-chain triacylglycerols and increases shorter-chain ceramides, while also modulating other lipid droplet-related lipid classes in a chain-length-dependent manner. Collectively, these findings suggest that METTL7B regulates lipid metabolism and malignant behavior in pancreatic cancer and may represent a potential therapeutic target.

## Introduction

Altered cellular metabolism is a hallmark of cancer^1^. While glucose and glutamine metabolism have been extensively characterized, emerging evidence highlights a critical role for lipid metabolism in cancer progression^2,3^. In pancreatic cancer, dysregulated lipid metabolism and lipid accumulation have been increasingly linked to disease development^4–10^. Nevertheless, the mechanisms by which lipid accumulation promotes pancreatic cancer remain poorly understood.

Lipids that accumulate in cells are stored in lipid droplets (LDs), organelles that serve as hubs of lipid metabolism and play dynamic roles in lipid homeostasis, energy balance, and signaling^11–13^. LDs sequester excess lipids and regulate lipid availability and metabolic signaling pathways. Dysregulation of LD formation and turnover has been implicated in a range of diseases, including metabolic disorders and cancer^14,15^. In pancreatic cancer, accumulating evidence suggests that LDs may support cancer cell survival and proliferation^16,17^, although the precise contribution of LDs to tumor development and progression is still being elucidated.

Many biological processes are regulated by chemical modifications of DNA, RNA, proteins, and small molecules, among which methylation is particularly prominent^18^. Methylation is catalyzed by diverse methyltransferases with distinct substrate specificities^19,20^. Methylation is essential for a wide range of normal cellular functions, and its dysregulation can drive human diseases, including cancer^21–25^. Nevertheless, for many methyltransferases, both their physiological substrates and their contributions to tumor biology remain poorly defined. Among these, methyltransferases that act on non-canonical substrates such as thiols are less well characterized.

METTL7B (methyltransferase-like 7B; also known as thiol methyltransferase 1B, TMT1B) is an alkyl thiol methyltransferase that catalyzes the methylation of exogenous thiols. METTL7B, together with its paralog METTL7A (also known as TMT1A and AAM-B), was recently identified as a thiol methyltransferase that methylates exogenous thiol-containing compounds such as 7α-thiospironolactone and captopril^26,27^. Furthermore, Robey et al. demonstrated that METTL7B, as well as METTL7A, confers resistance to thiol-based histone deacetylase inhibitors, such as romidepsin^28^. Beyond its role in thiol methylation, METTL7B has been implicated in the progression of multiple cancers. METTL7B is reported to be upregulated in several tumor types, including breast cancer^29^, thyroid cancer^30,31^, non-small cell lung cancer (NSCLC)^32^, clear cell renal cell carcinoma (ccRCC)^33^, lung adenocarcinoma^34,35^, and glioma^36–38^. Despite these observations, the mechanisms underlying METTL7B upregulation remain largely unknown.

METTL7B exhibits context-dependent subcellular localization. In humans, mice, and rats, METTL7B localizes to the endoplasmic reticulum (ER) under unstimulated conditions^29,39,40^. When intracellular LDs increase—for example, after partial hepatectomy or oleic acid treatment—METTL7B translocates from the ER to the LD surface^39–41^. This dynamic localization pattern suggests that METTL7B may participate in the biogenesis and function of LDs. However, the functional significance of its recruitment to LDs and its contribution to lipid metabolism and tumor progression in pancreatic cancer remain unclear.

Here, we demonstrate that METTL7B is upregulated in pancreatic cancer and that high METTL7B expression is associated with poor patient prognosis. Using loss-of-function approaches, we show that METTL7B is required for pancreatic cancer cell proliferation, migration, and metastasis. We also identify HNF4A and HNF4G, well-established regulators of lipid homeostasis, as transcription factors that contribute to METTL7B upregulation in pancreatic cancer. Furthermore, we demonstrate that METTL7B localizes around LDs and promotes their accumulation in pancreatic cancer cells. Lipidomic analyses reveal that METTL7B depletion selectively reduces long-chain triacylglycerols (TGs) and increases shorter-chain ceramides, while also remodeling other lipid droplet-related lipid classes in a chain-length-dependent manner. Collectively, these findings indicate that METTL7B contributes to lipid metabolism and malignant progression in pancreatic cancer and may serve as a potential therapeutic target.

## Results

### Pan-cancer profiling of methyltransferases identifies METTL7B as a prognostic factor in pancreatic cancer

To identify methyltransferases with potential clinical relevance across various cancers, we analyzed the expression and prognostic significance of 155 methyltransferases using the TCGA and GTEx datasets. Many methyltransferases were dysregulated across cancers, with prominent alterations in pancreatic cancer, one of the most lethal malignancies^42^ (Fig. 1a, S1a). However, most showed no significant association with patient survival (Fig. 1b, S1b).

**Figure 1.**
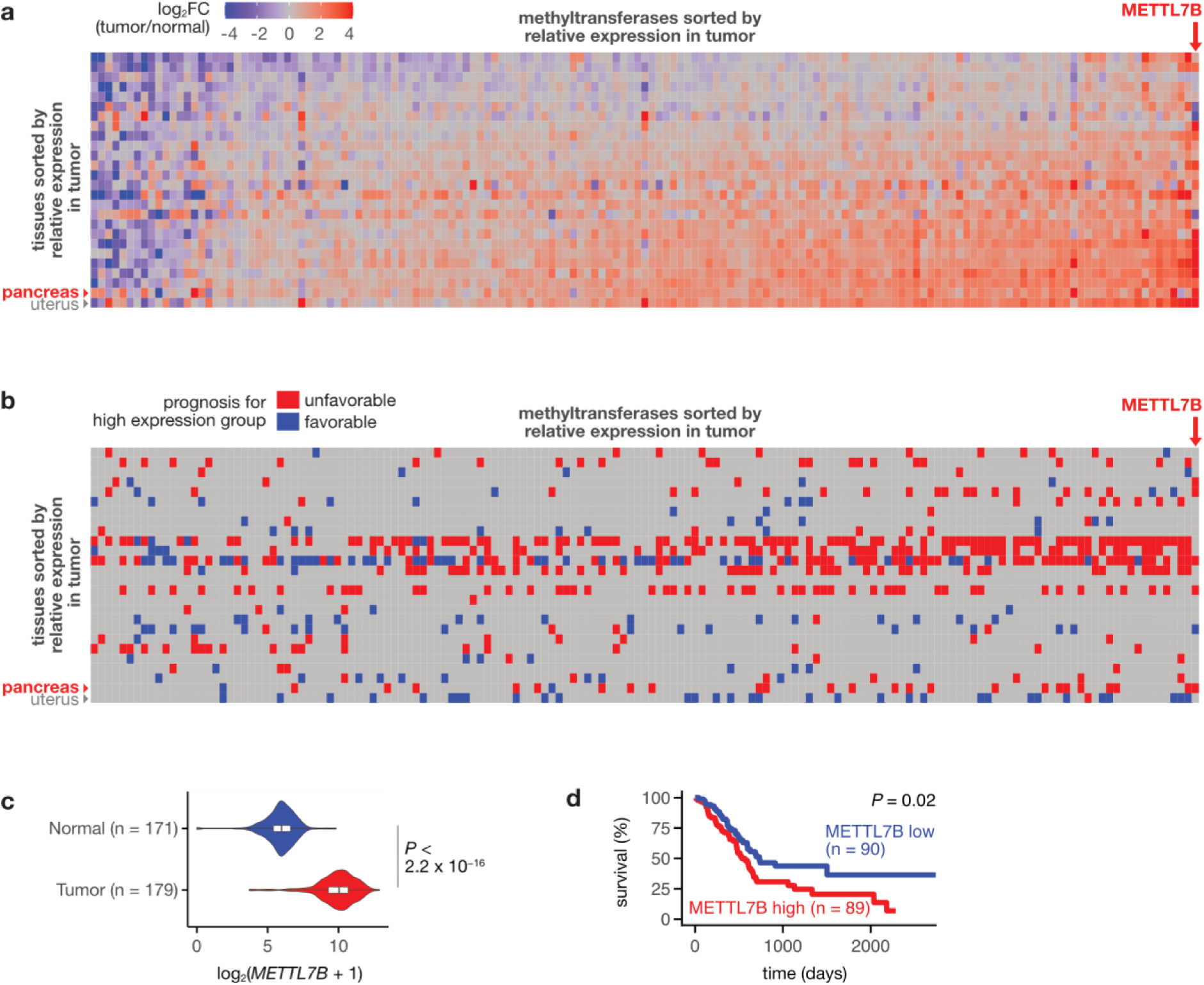
Pan-cancer profiling of methyltransferases identifies METTL7B as a prognostic factor in pancreatic cancer. **a** Heatmap showing log_2_ fold change (tumor vs. normal) for 155 methyltransferases across multiple cancer types. *METTL7B* and pancreas are indicated by an arrow. Methyltransferases and cancer types are sorted by relative expression in tumor versus normal tissue. **b** Heatmap showing the prognostic association of high methyltransferase expression with overall survival across multiple cancer types. Red indicates unfavorable prognosis, blue indicates favorable prognosis, and gray indicates no significant association. *METTL7B* and pancreas are indicated by an arrow. **c** Violin plot showing *METTL7B* expression in normal and tumor pancreatic tissues (normal, n = 171; tumor, n = 179). Wilcoxon test. **d** Kaplan–Meier survival curve for pancreatic cancer patients stratified by *METTL7B* expression. The red and blue lines represent patients with high and low *METTL7B* expression, respectively; n (high) = 89, n (low) = 90.

Among these genes, METTL7B was one of the most consistently upregulated methyltransferases (Fig. 1a, S1a). Consistent with previous reports^30,32,34–37,43–45^, METTL7B was upregulated in multiple cancers, including papillary thyroid cancer, lung adenocarcinoma, and esophageal adenocarcinoma (Fig. S1a). Notably, pancreatic cancer showed significant METTL7B upregulation (Fig. 1a, 1c, S1a). Moreover, a clear association between high METTL7B expression and poor patient prognosis was observed in pancreatic cancer (Fig. 1b, 1d, S1b). Together, these results indicate that METTL7B upregulation correlates with poor clinical outcomes in pancreatic cancer.

### METTL7B is required for pancreatic cancer cell proliferation, migration, and metastasis

To investigate the role of METTL7B in pancreatic cancer pathogenesis, we established two independent METTL7B-knockout (KO) AsPC-1 cell clones using the CRISPR–Cas9 system (Fig. 2a, b). Using an intrasplenic injection model of pancreatic cancer metastasis^46,47^, we assessed the impact of METTL7B depletion on metastatic potential *in vivo* (Fig. 2c). METTL7B-KO cells showed markedly reduced metastasis compared with parental AsPC-1 cells (Fig. 2c-e). Consistently, METTL7B-KO clones exhibited impaired migration and decreased proliferation compared with parental AsPC-1 cells *in vitro* (Fig. 2f-h). Together, these findings indicate that METTL7B is required for pancreatic cancer cell proliferation, migration, and metastasis.

**Figure 2.**
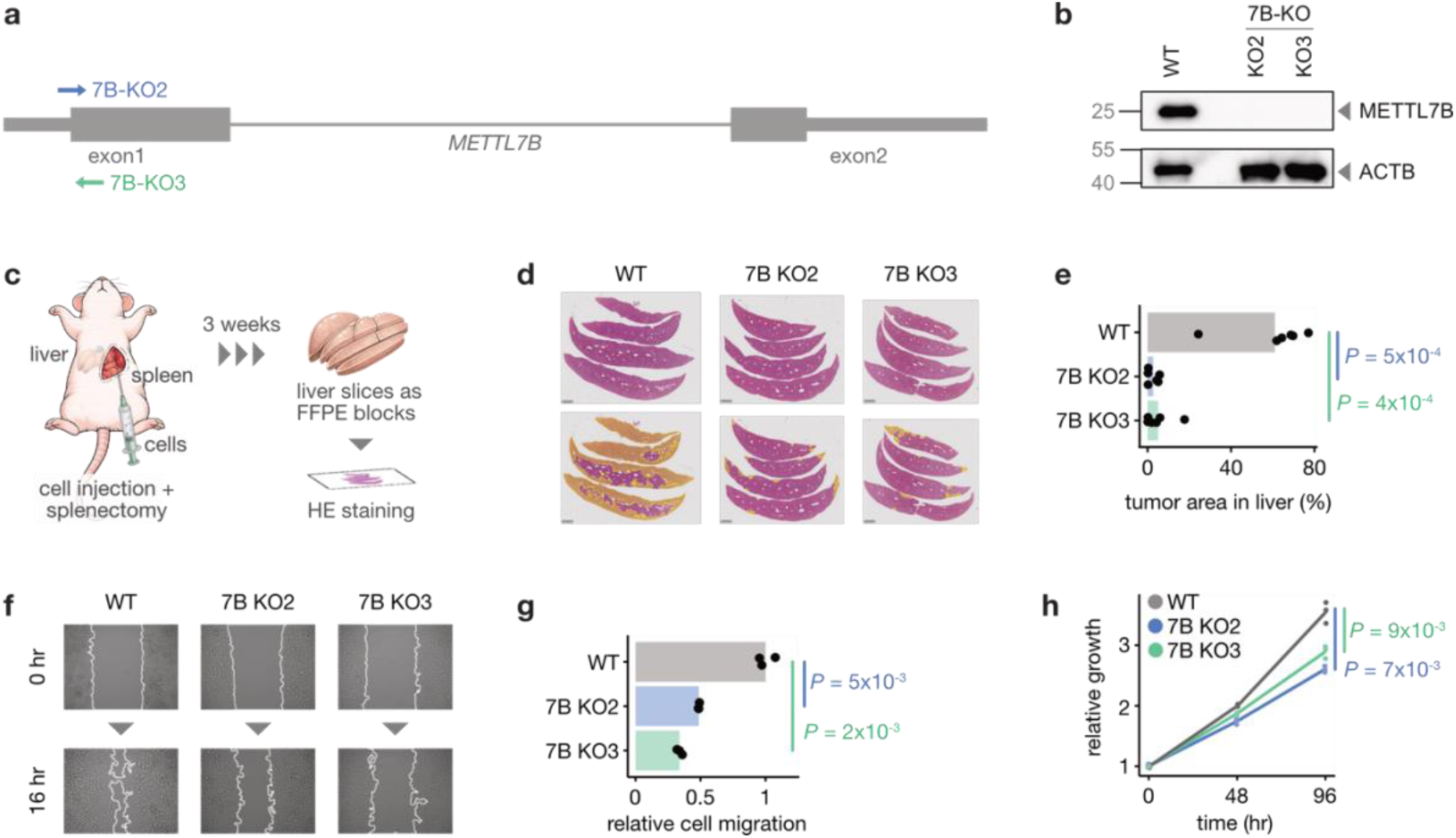
METTL7B knockout impairs growth, migration and metastasis of AsPC-1 cells. **a** Schematic of the *METTL7B* locus showing the positions of CRISPR–Cas9 guide RNAs (7B-KO2 and 7B-KO3 indicated by arrows) designed near the start codon of the METTL7B open reading frame (ORF). Coding regions of the ORF are depicted as thick rectangles, exons as thin rectangles, and introns as very thin rectangles. **b** Western blotting analysis of METTL7B protein levels in parental AsPC-1 cells (WT) and two independent METTL7B-KO clones, with ACTB used as a loading control. **c** Schematic workflow for the intrasplenic liver metastasis assay. AsPC-1 clone cells were injected into the spleen, and splenectomy was performed immediately afterward. Mice were maintained for 3 weeks, then livers were harvested and processed into FFPE blocks. Sections were stained with hematoxylin and eosin (H&E) for histological analysis. **d** Representative histological images of liver metastases formed by WT, 7B KO2, and 7B KO3 cells. Upper panels show low-magnification H&E–stained images of liver sections from mice injected with the indicated AsPC-1 clones. Lower panels show the corresponding images with tumor regions annotated (yellow). Scale bar, 2 mm. **e** Quantification of liver tumor area in the metastasis assay for WT, 7B KO2, and 7B KO3 cells. Each dot represents an individual mouse. Student’s *t*-test. n = 6 mice per group. **f** Representative brightfield images from wound healing assays comparing the migratory capabilities of WT, 7B KO2, and 7B KO3 cells. Images are shown at 0 hours (immediately after scratch formation) and after 16 hours. **g** Quantification of relative cell migration of WT, 7B KO2, and 7B KO3 cells, with WT migration normalized to 1.0. Individual data points from replicates are shown. Student’s *t*-test. **h** Relative growth curves of WT cells and two independent METTL7B-knockout clones (7B KO2, and 7B KO3) over 96 hours. All cell lines are normalized to a relative growth of 1 at 0 hours. Student’s *t*-test at 96 hours.

To investigate how METTL7B regulates pancreatic cancer cell proliferation and migration, we performed RNA-seq analysis using the two independent METTL7B-KO AsPC-1 clones. The two METTL7B-KO clones showed highly similar expression changes (Fig. 3a), with 948 genes downregulated and 1,057 genes upregulated upon METTL7B depletion (Fig. 3b, Supplementary Table S1). GO analysis revealed that the upregulated and downregulated genes were associated with 243 and 242 biological processes, respectively, corresponding to 420 unique processes in total. These processes were further classified into 30 clusters based on semantic similarity (Fig. 3c, Supplementary Table S2). Consistent with the phenotypes observed upon METTL7B depletion, enriched processes included those related to cell motility and proliferation (cluster 21). In addition, we identified processes related to organelle organization (cluster 24) and lipid metabolism (cluster 9) (Fig. 3c). Upregulated genes were primarily associated with responses to stimuli (cluster 1 in Fig. 3c; Fig. 3d), whereas downregulated genes were enriched for cell cycle and DNA replication processes (clusters 2 and 27 in Fig. 3c; Fig. 3e), suggesting that METTL7B regulates pancreatic cancer cell proliferation, at least in part, via these pathways.

**Figure 3.**
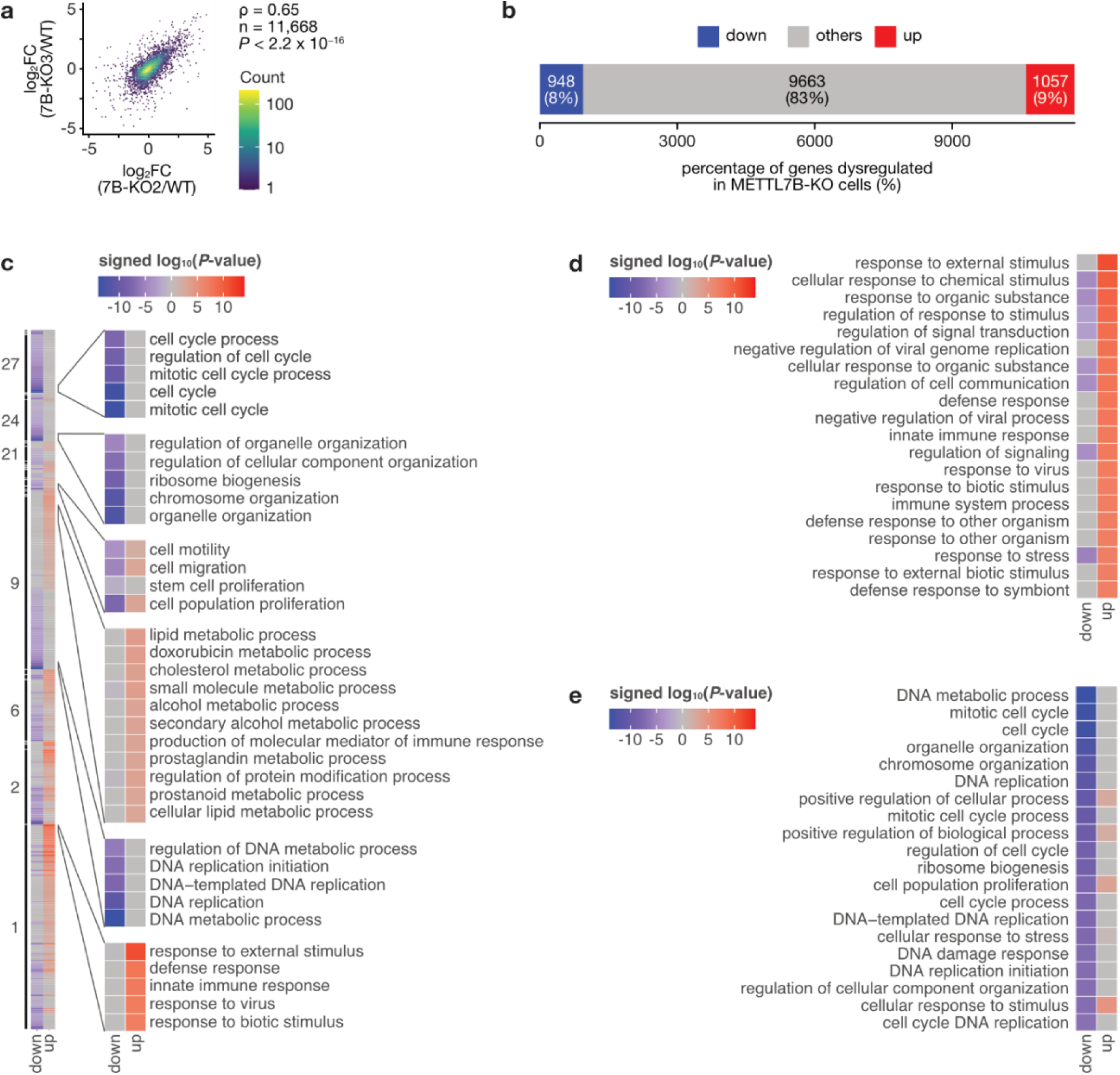
METTL7B knockout alters biological processes linked to proliferation and lipid metabolism. **a** Scatter plot showing the correlation of log_2_ fold changes (KO vs. WT) for each gene between the two independent METTL7B-KO clones (7B KO2 and 7B KO3). Spearman’s correlation test. **b** Stacked bar chart showing the number and percentage of genes that are downregulated, unchanged, or upregulated in METTL7B-KO cells relative to parental cells. **c** Heatmap showing the enrichment of Gene Ontology (GO) biological processes among genes upregulated and downregulated in METTL7B-KO cells. Enriched processes were grouped into 30 clusters based on semantic similarity and within each cluster, sorted by enrichment strength. Colors indicate the signed log_10_(*P*-value), with positive values for processes enriched in upregulated genes and negative values for processes enriched in downregulated genes. **d** Heatmap of the top 20 GO biological processes most strongly enriched among genes downregulated in METTL7B-KO cells, ranked by signed log_10_(*P*-value). **e** Heatmap of the top 20 GO biological processes most strongly enriched among genes upregulated in METTL7B-KO cells, ranked by signed log_10_(*P*-value).

### HNF4A and HNF4G contribute to METTL7B upregulation in pancreatic cancer

To elucidate the mechanism underlying METTL7B upregulation in pancreatic cancer, we searched for transcription factors (TFs) that bind near the METTL7B transcription start site (TSS) and whose expression correlates with METTL7B in pancreatic cancer. Integration of ChIP-Atlas binding data with TCGA expression data identified hepatocyte nuclear factor 4α (HNF4A) and hepatocyte nuclear factor 4γ (HNF4G) as candidate regulators of METTL7B expression in pancreatic cancer (Fig. 4a, 4b).

**Figure 4.**
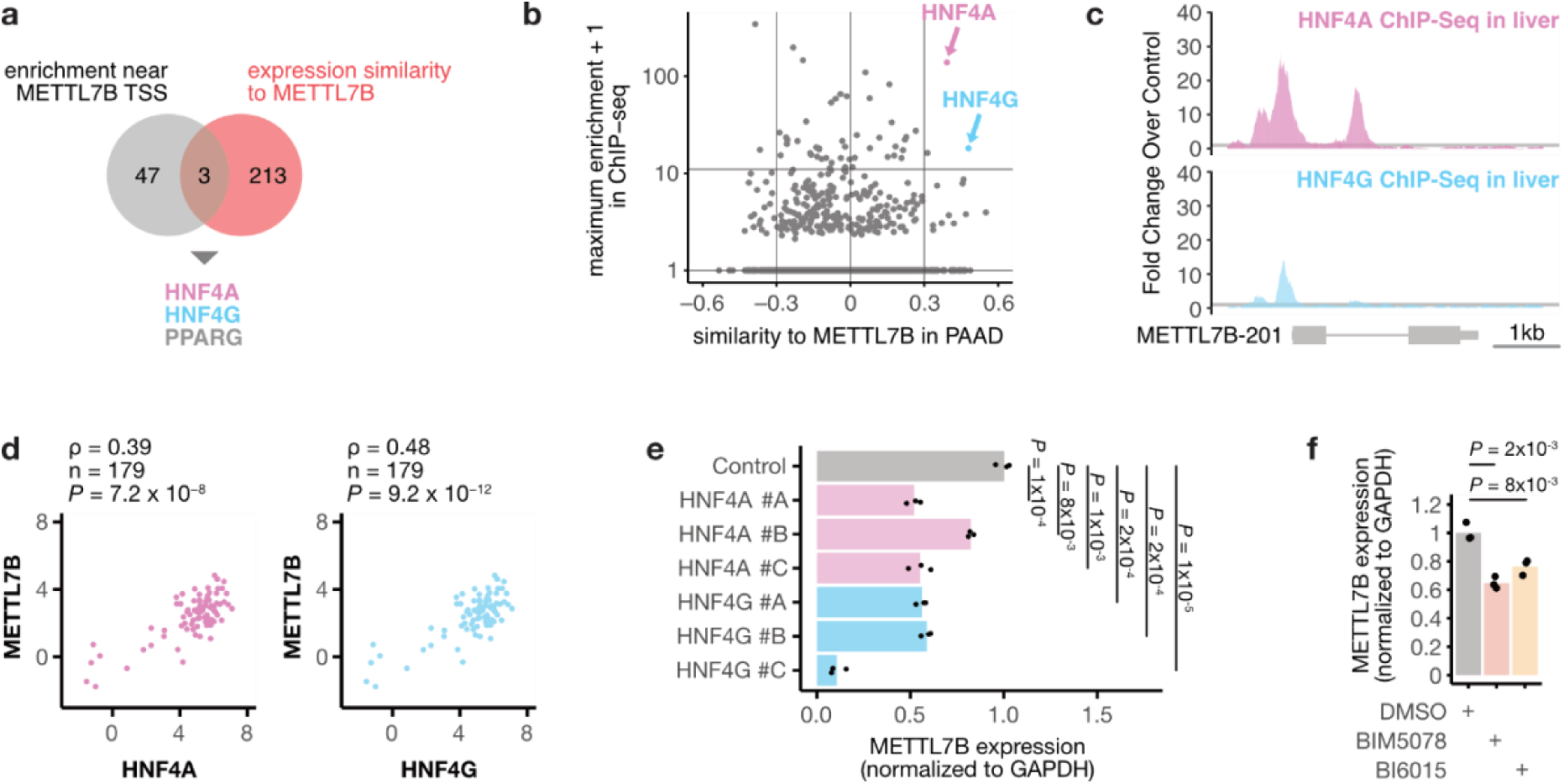
HNF4A and HNF4G are transcriptional regulators of METTL7B expression in pancreatic cancer. **a** Venn diagram showing the overlap between transcription factors (TFs) enriched near the *METTL7B* transcription start site (TSS) in ChIP-Atlas (maximum enrichment score > 10) and TFs whose expression is positively correlated with *METTL7B* in TCGA pancreatic adenocarcinoma (PAAD). HNF4A and HNF4G are among the TFs in the overlapping region. **b** Scatter plot of TFs showing expression similarity to *METTL7B* in TCGA PAAD (x-axis; correlation coefficient) versus enrichment near the *METTL7B* TSS in public ChIP-seq datasets (y-axis; maximum enrichment score), highlighting HNF4A and HNF4G. **c** Coverage tracks showing enrichment of HNF4A and HNF4G ChIP-seq reads near the *METTL7B* TSS in liver-derived samples. **d** Scatter plots showing correlations between *METTL7B* expression and *HNF4A* (left) or *HNF4G* (right) expression in TCGA PAAD tumors. n = 179. Spearman’s correlation test. **e** Relative *METTL7B* mRNA expression in AsPC-1 cells following knockdown of *HNF4A* or *HNF4G*, normalized to *GAPDH*. Individual data points represent biological replicates. Student’s *t*-test. **f** Relative *METTL7B* mRNA expression in AsPC-1 cells treated with HNF4 inhibitors (BIM5078 or BI6015) or DMSO control, normalized to *GAPDH*. Individual data points represent biological replicates. Student’s *t*-test.

HNF4A and HNF4G are members of the HNF4 family, which belongs to the nuclear receptor superfamily^48^ and is known to play essential roles in lipid homeostasis^49–52^. Public ChIP-seq data confirmed that both HNF4A and HNF4G bind near the METTL7B TSS in liver-derived cells, with HNF4A binding also detected in pancreas-derived tissues (Fig. 4b, 4c, S2a, S2b). In pancreatic cancer tissues, HNF4A and HNF4G levels strongly correlated with METTL7B expression in both TCGA and Australian Pancreatic Cancer Genome Initiative (APGI) datasets (Fig. 4b, 4d, S2c). Moreover, similar to METTL7B, both HNF4A and HNF4G were upregulated in pancreatic cancer tissues compared with normal tissues (Fig. S2d).

We then tested this regulatory relationship. Knockdown of HNF4A or HNF4G reduced METTL7B expression (Fig. 4e, S2e, S2f). Pharmacological inhibition of HNF4A with BIM5078^53^ and BI6015^53^ also reduced METTL7B expression (Fig. 4f). Together, these findings indicate that HNF4A and HNF4G contribute to METTL7B upregulation in pancreatic cancer.

### METTL7B localizes around and promotes the accumulation of lipid droplets

To investigate the function of METTL7B in pancreatic cancer cells, we examined its subcellular localization. While initial proteomic analyses reported METTL7B localization to the Golgi apparatus^54^, subsequent immunostaining studies indicated its localization to the endoplasmic reticulum (ER) and lipid droplets (LDs)^29^. In our immunostaining analysis, METTL7B formed numerous circular structures (Fig. 5a). Co-staining with organelle markers revealed that these structures, which surrounded LDs, were in contact with the ER and mitochondria (Fig. 5a).

**Figure 5.**
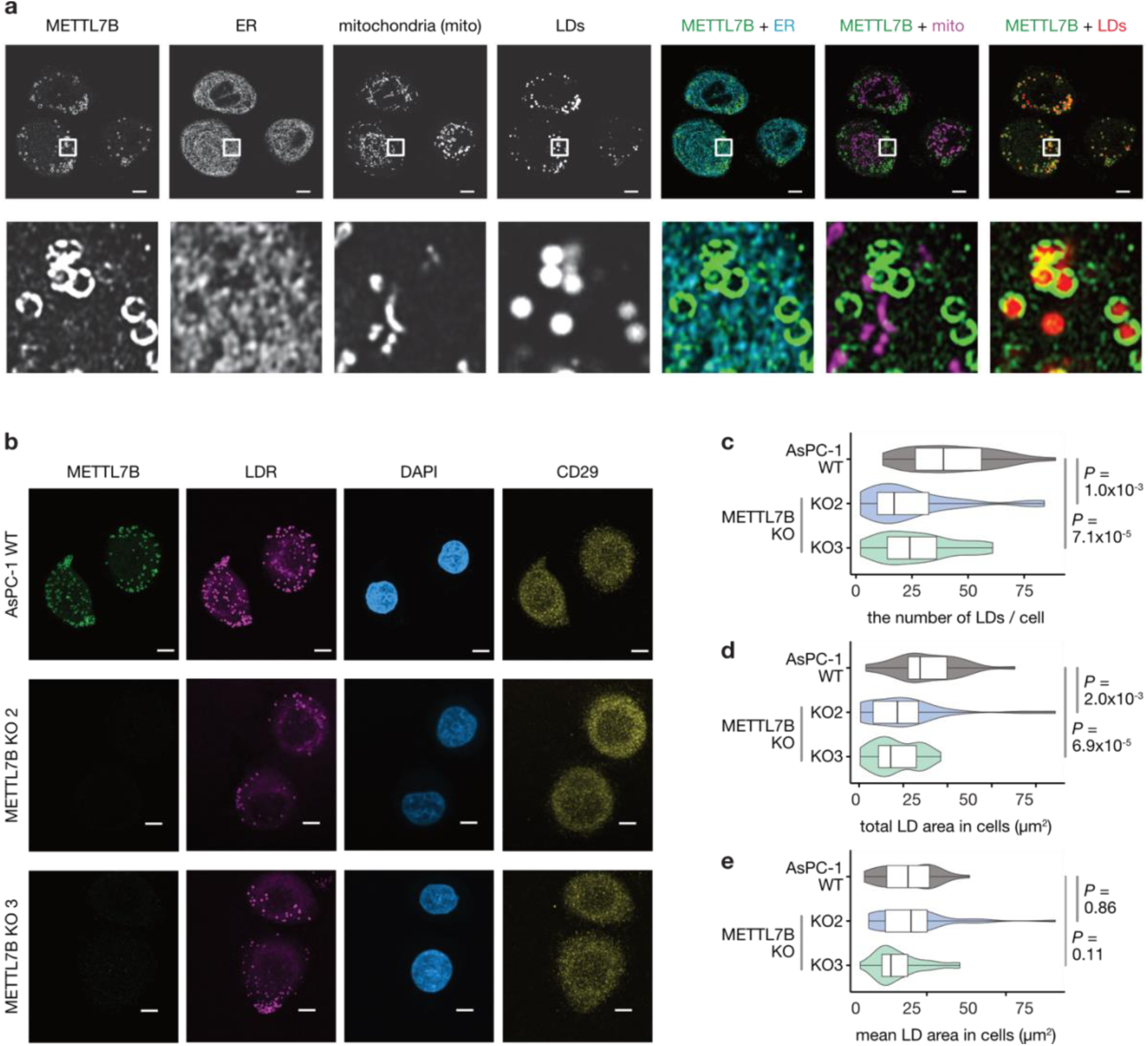
METTL7B localizes around and promotes the accumulation of lipid droplets in pancreatic cancer cells. **a** Confocal microscopic images of AsPC-1 cells stained for METTL7B (anti-METTL7B; Alexa Fluor 488, green), endoplasmic reticulum (ER; anti-PDIA3; Alexa Fluor 647, magenta), mitochondria (MitoTracker Red CMX Ros, red) and lipid droplets (LDs; Lipi-Blue, blue). Insets show higher-magnification views highlighting METTL7B-positive structures surrounding LDs and contacting ER and mitochondria. Scale bar: 5 µm. **b** Confocal microscopic images of parental (wild-type, WT) AsPC-1 cells and METTL7B-knockout (KO) clones (7B KO2 and 7B KO3) stained for METTL7B (anti-METTL7B; Alexa Fluor 488, green) and integrin β1 (CD29; anti-CD29; Alexa Fluor 555, orange), with LDs labeled with Lipi-DeepRed (magenta) and nuclei counterstained with DAPI (blue). CD29 staining was used to define cell regions. Scale bar: 5 µm. **c** Violin plots showing the distribution of the number of LDs per cell in WT AsPC-1 and METTL7B KO cells. Wilcoxon test. WT n = 31, KO2 n = 35, KO3 n = 33. **d** Violin plots showing the distribution of total LD area per cell in WT AsPC-1 cells and METTL7B KO cells. Wilcoxon test. WT n = 31, KO2 n = 35, KO3 n = 33. **e** Violin plots showing the distribution of mean LD area per cell in WT AsPC-1 cells and METTL7B KO cells. Wilcoxon test. WT n = 31, KO2 n = 35, KO3 n = 33.

Previous studies have shown that METTL7B predominantly localizes around LDs under lipid-rich conditions^39–41^, yet its function at LDs remains unclear. We therefore examined whether METTL7B regulates LD abundance in pancreatic cancer cells. LD staining showed that METTL7B-KO clones exhibited a significant reduction in both LD number and total LD area, whereas the average size of individual LDs remained unchanged compared with parental AsPC-1 cells (Fig. 5b–e). Taken together, these results demonstrate that METTL7B localizes around LDs and promotes their accumulation in pancreatic cancer cells.

### METTL7B upregulates triacylglycerols and downregulates ceramides

Multiple lines of evidence suggested that METTL7B regulates lipid metabolism in pancreatic cancer cells. RNA-seq analysis of METTL7B-KO cells showed enrichment of genes involved in lipid metabolic processes (cluster 9 in Fig. 3c). In addition, HNF4A and HNF4G, identified as upstream regulators of METTL7B (Fig. 4), are well-established regulators of lipid homeostasis^49,50,55–57^. Moreover, METTL7B localizes around LDs and promotes their accumulation in pancreatic cancer cells (Fig. 5). These results prompted us to investigate the specific lipid species affected by METTL7B depletion.

Lipidomic and metabolomic analyses identified 4,998 peaks, of which 181 were downregulated and 408 were upregulated in METTL7B-KO cells (Fig. 6a, S3a, Supplementary Table S3). Dysregulated peaks were more prominent in lipidomic than metabolomic profiles (Fig. 6b). Peak annotation using the LIPID MAPS database revealed that downregulated peaks were enriched for glycerolipids [GL], especially triradylglycerols [GL03], whereas upregulated peaks were enriched for sphingolipids [SP], particularly ceramides (Fig. 6c, S3b). Notably, these lipid changes were consistent with the observed phenotypes of METTL7B-KO cells: triradylglycerols constitute the hydrophobic core of LDs, whereas ceramides are sphingolipids that regulate apoptosis and proliferation, acting as tumor suppressors^58,59^.

**Figure 6.**
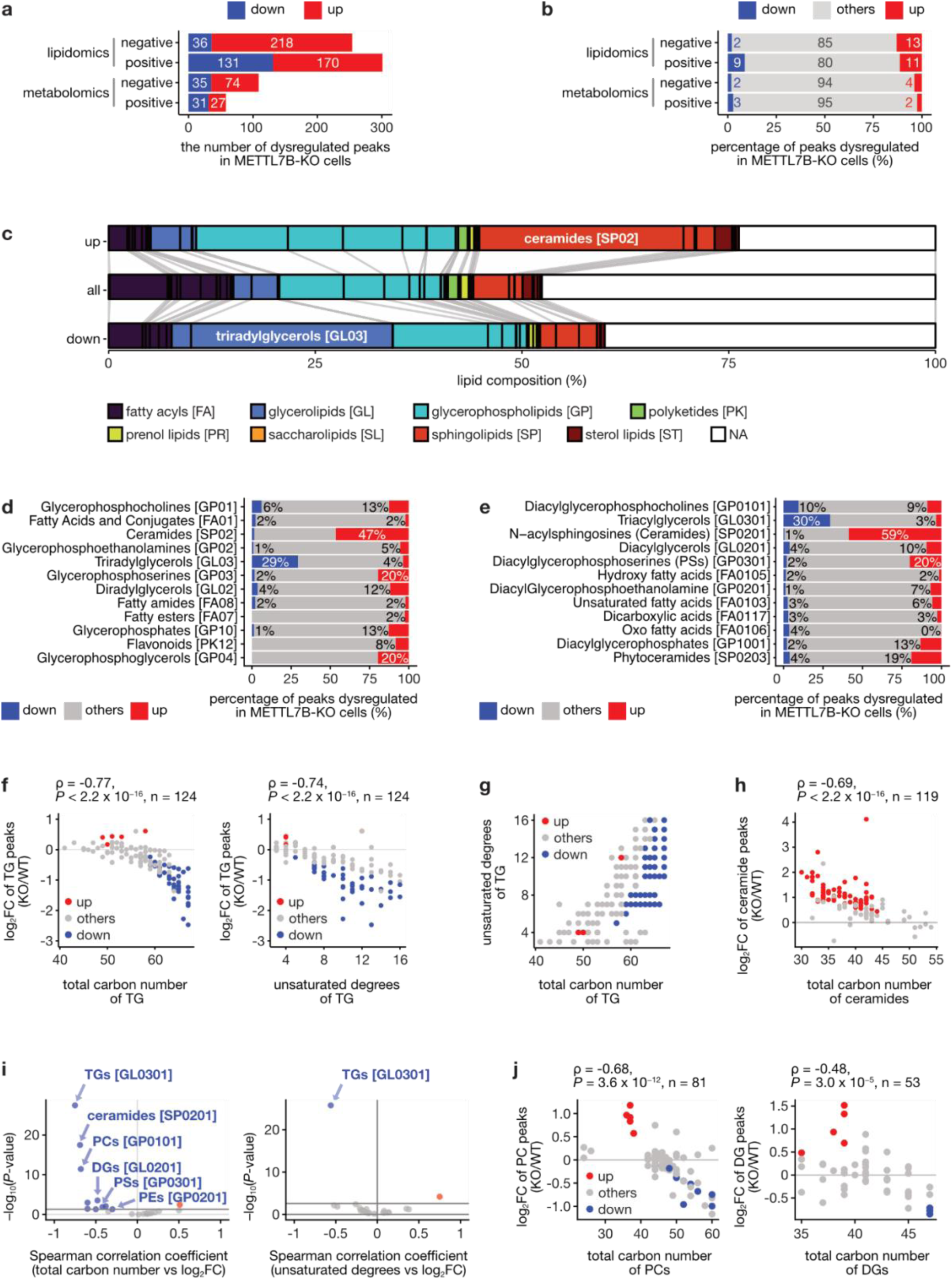
METTL7B knockout remodels lipidomic profiles. **a** Stacked bar chart showing the total number of significantly dysregulated peaks in METTL7B-KO cells. Peaks are categorized by detection method (lipidomics or metabolomics), ion mode (negative or positive), and direction of regulation (downregulated in blue, upregulated in red). **b** Stacked bar chart showing the percentage of significantly dysregulated peaks in METTL7B-KO cells, using the same categorization as in panel a. **c** Alluvial plot illustrating the lipid composition of dysregulated peaks. The top bar represents upregulated peaks, the middle bar represents all peaks, and the bottom bar represents downregulated peaks. The bars are segmented and colored by lipid class. **d** Stacked bar charts showing the percentage of dysregulated peaks within each main lipid classes. Only classes with >100 detected peaks are shown. **e** Stacked bar charts showing the percentage of dysregulated peaks within each lipid subclass. Only subclasses with >100 detected peaks are shown. **f** Scatter plots showing the log₂ fold change (KO/WT) of triacylglycerol (TG) peaks versus their total carbon number (left) and degree of unsaturation (right). Points are colored by regulation status (red: upregulated, blue: downregulated, grey: others). Spearman’s correlation test. **g** Scatter plot with individual data points showing the distribution of total carbon numbers (left) and degrees of unsaturation (right) of TGs for upregulated (red), others (grey), and downregulated (blue) peaks. Wilcoxon test. **h** Scatter plot showing the log₂ fold change (KO/WT) of ceramide peaks against their total carbon number. Points are colored by regulation status (red: upregulated, blue: downregulated, grey: others). Spearman’s correlation test. **i** Scatter plots showing the Spearman correlation coefficient versus −log_10_(*P*) for the relationship between lipid characteristics and log_2_ fold change. Left panel: correlation between total carbon number and log_2_FC for various lipid classes. Right panel: correlation between degrees of unsaturation and log_2_FC for various lipid classes. **j** Scatter plots showing the log_2_ fold change (KO/WT) of phosphatidylcholine (PC) peaks (left) and diacylglycerol (DG) peaks (right) versus their total carbon number. Points are colored by regulation status (red: upregulated, blue: downregulated, grey: others). Spearman’s correlation test.

To quantify the extent of dysregulation across different lipid classes, we calculated the percentage of significantly altered peaks in each class. Among the major lipid classes, ceramides [SP02] showed the highest percentage of upregulated peaks (47%), whereas triradylglycerols [GL03] showed a substantial percentage of downregulated peaks (29%) (Fig. 6d, S3c). Lipid sub-class analysis further strengthened this pattern: 59% of ceramide [SP0201] peaks were upregulated, and 30% of triacylglycerol [GL0301] peaks were downregulated in METTL7B-KO cells (Fig. 6e, S3d). Ceramides [SP02] mainly comprised ceramides [SP0201], dihydroceramides [SP0202], and phytoceramides [SP0203], with the most pronounced upregulation observed in ceramides [SP0201] (Fig. S4a). Most triradylglycerols [GL03] were assigned to triacylglycerols (TGs) [GL0301] (Fig. S4b). Together, these data indicate that METTL7B depletion shifts the lipid profile toward higher ceramide levels and lower TG levels.

Further analysis revealed that TG reduction was selective. The extent of reduction showed a negative correlation with both the total acyl chain length and degrees of unsaturation (Fig. 6f). Downregulated TG species upon METTL7B knockout displayed longer acyl chains and higher unsaturation levels than other TG species (Fig. S3c). The distribution of TG species confirmed that downregulated TGs were enriched among those with higher total carbon numbers, not higher degrees of unsaturation (Fig. 6g). These results indicate that METTL7B preferentially regulates the abundance of long-chain TGs.

To determine whether ceramide upregulation showed similar structural specificity, we analyzed the relationship between ceramide structural features and the extent of upregulation upon METTL7B KO. Ceramides exist in multiple forms with different acyl chain lengths and degrees of unsaturation, and recent studies have revealed that these structural variations are associated with distinct functional roles^60^. Ceramides with shorter acyl chains showed a selective increase in METTL7B KO cells (Fig. 6h, S4d), whereas no clear correlation was observed with degrees of unsaturation (Fig. S4e). These results indicate that METTL7B depletion is associated with chain-length-dependent remodeling of ceramide species.

Because both TGs and ceramides showed chain-length-dependent relationships between total carbon numbers and peak intensity changes, we extended the analyses to other lipid classes. Similar but weaker negative correlations were observed in multiple lipid classes, including phosphatidylcholines (PCs, diacylphosphatidylcholines) [GP0101], diacylglycerols (DGs) [GL0201], phosphatidylserines (PSs, diacylglycerophosphoserines) [GP0301], and phosphatidylethanolamines (PEs, diacylglycerophosphoethanolamines) [GP0201] (Fig. 6i, 6j, S4f). In contrast, this pattern was not observed between log₂FC and degrees of unsaturation for these lipid classes (Fig. 6i). PCs are the main component of the phospholipid monolayer of LDs^61–63^, and DGs, PSs, PEs, and ceramides are also components of the LD lipidome^63^. Together, these results indicate that METTL7B depletion is associated with chain-length-dependent changes across multiple LD-associated lipid classes.

## Discussion

In this study, we investigated the role and regulation of METTL7B in pancreatic cancer. Analysis of large-scale transcriptomic datasets revealed that METTL7B is upregulated in pancreatic cancer tissues, and elevated expression is associated with poor patient prognosis, consistent with previous reports in other cancer types. We further demonstrated that METTL7B is required for the proliferation, migration, and metastasis of pancreatic cancer cells. Mechanistically, we identified HNF4A and HNF4G, well-established regulators of lipid metabolism, as transcription factors that drive METTL7B upregulation in pancreatic cancer. We showed that METTL7B localizes around lipid droplets and promotes their accumulation, and that loss of METTL7B leads to marked changes in lipid metabolism, including decreased triacylglycerols and increased ceramides. These findings establish METTL7B as a regulator of lipid metabolism and tumor progression in pancreatic cancer, suggesting that targeting the METTL7B pathway may represent a promising therapeutic strategy.

A key finding of this study is that METTL7B promotes LD accumulation and is associated with lipid remodeling in pancreatic cancer cells. Dysregulated lipid metabolism and lipid accumulation are increasingly recognized as drivers of pancreatic cancer^2–10,64^, yet the mechanisms linking these features to tumor progression remain unclear. METTL7B has previously been reported to localize around LDs when intracellular LDs are induced^39,40^; however, its function at LDs has remained unclear. In this study, we demonstrated that METTL7B localizes around LDs in pancreatic cancer cells under basal conditions (i.e., without oleic acid [OA] treatment), extending previous observations to unstimulated conditions. Furthermore, depletion of METTL7B led to a decrease in both the number and total area of LDs, indicating that METTL7B regulates LD abundance.

Lipidomic analysis revealed that this reduction in LD abundance is associated with marked changes in lipid composition. Triacylglycerols—one of the main components of LDs—were significantly reduced in METTL7B-KO cells. Notably, this reduction was not uniform across all TG species; rather, METTL7B depletion selectively decreased long-chain triacylglycerols, while shorter-chain TGs were largely unaffected. This specificity suggests that METTL7B may preferentially influence the synthesis, storage, or turnover of specific TG species that contribute to LD accumulation. In addition to the reduction in TGs, METTL7B depletion led to an increase in ceramides, particularly those with shorter acyl chains. Given that ceramides can act as tumor suppressors and promote apoptosis^58^, this increase may contribute to the impaired proliferation observed in METTL7B-deficient cells. The opposing effects of METTL7B on TGs and ceramides—promoting accumulation of long-chain TGs while suppressing shorter-chain ceramides—suggest that METTL7B coordinates lipid metabolism to maintain a lipid profile that supports cancer cell proliferation, migration, and metastasis. Beyond TGs and ceramides, METTL7B also modulates other LD-related lipid classes, including PCs, DGs, PSs, and PEs, in a chain-length-dependent manner. Since PCs are the main component of the phospholipid monolayer of LDs^63,65^, and DGs, PSs, PEs, and ceramides are components of LDs^63^, these data support a model in which METTL7B regulates both the quantity and quality of LDs. ^15,66^

A key strength of this study is the identification of HNF4A and HNF4G as transcription factors that contribute to METTL7B upregulation in pancreatic cancer. Although previous studies have shown that METTL7B is upregulated in several cancers—including thyroid cancer^30^, NSCLC^32^, ccRCC^33^, lung adenocarcinoma^34,35^, and glioma^36,37,44^—compared with their normal tissues, the mechanisms underlying METTL7B upregulation have remained unclear. In this study, we identified HNF4A and HNF4G as transcription factors that drive the upregulation of METTL7B in pancreatic cancer using two complementary analyses of public datasets: (1) co-expression with METTL7B in pancreatic cancer tissue in the TCGA dataset, and (2) binding near the transcription start site (TSS) of METTL7B in public ChIP-seq datasets. Consistent with these predictions, knockdown of HNF4A or HNF4G and pharmacological inhibition of HNF4A reduced METTL7B expression.

Notably, HNF4A and HNF4G are well-established regulators of lipid homeostasis with distinct roles in the liver and intestine. HNF4A plays critical roles in hepatic lipid homeostasis^49,50,52,56^, as shown by liver-specific deletion models in which loss of HNF4A causes marked disruptions in lipid metabolism, including decreased serum cholesterol and triglyceride levels, and reduced expression of genes involved in lipid transport and metabolism^52^. In the intestine, both HNF4A and HNF4G regulate lipid absorption, transport, and metabolism. These paralogs act redundantly to activate enhancer chromatin and upregulate the majority of transcripts enriched in the differentiated epithelium, including genes involved in lipid metabolism. When both HNF4 paralogs are deleted, cells fail to differentiate into enterocytes, the primary absorptive cells responsible for lipid uptake, processing, and metabolism in the intestine^66^. Beyond their role in enterocyte differentiation, HNF4A and HNF4G are required for fatty acid oxidation and renewal of intestinal stem cells^51^. Specifically, loss of HNF4A and HNF4G in intestinal organoids leads to compromised β-oxidation contribution to the TCA cycle metabolites and elevated fatty acid synthesis, indicating a critical role in regulating fatty acid oxidation pathways^51^. Furthermore, HNF4G deletion leads to intestinal lipid malabsorption and protection against diet-induced obesity^55^, while HNF4A deletion causes resistance to diet-induced obesity and influences whole-body energy expenditure^57^. The identification of HNF4A and HNF4G as regulators of METTL7B expression, combined with our finding that METTL7B regulates LD accumulation and lipid metabolism in pancreatic cancer cells, suggests that HNF4A/HNF4G may control lipid metabolism in pancreatic cancer, at least in part, through METTL7B.

Although METTL7B has been shown to methylate exogenous thiols such as captopril^26^, methanethiol^67^, and thiol-based histone deacetylase inhibitors^28^, its endogenous methylation target(s) remain unknown. Despite METTL7B’s demonstrated ability to methylate thiols, our lipidomic and metabolomic analyses did not detect altered levels of methylated thiols in response to METTL7B KO. We searched for potential endogenous methylation products by requiring that peaks (1) decreased in METTL7B-KO cells, (2) had a corresponding peak with m/z lower by one CH₂ unit, and (3) contained sulfur in their formula; no such peaks were identified. The ability of previous studies to detect METTL7B-mediated methylation likely reflects their use of purified recombinant METTL7B with specific exogenous substrates under controlled conditions, where substrates could be added fresh and protected from oxidation, and detection methods were optimized for those particular compounds. In contrast, our analysis of complex biological samples may have failed to detect endogenous methylated thiols due to technical challenges, including thiol oxidation during sample preparation and analysis^68,69^, as well as low concentrations or rapid metabolism of endogenous methylated thiols. Further research using targeted approaches with optimized sample preparation and detection methods will be required to identify potential endogenous methylation targets and clarify their association with METTL7B-mediated cancer progression.

To understand how METTL7B promotes pancreatic cancer progression, we analyzed the functional consequences of METTL7B depletion. We demonstrated that METTL7B is required for pancreatic cancer cell proliferation, migration, and metastasis. RNA-seq analysis in METTL7B-KO cells revealed that cell cycle and DNA replication were among the most significantly affected processes, suggesting that METTL7B is required for these processes in pancreatic cancer cells. The role of METTL7B in regulating the cell cycle appears to be conserved across multiple cancer types: METTL7B deletion downregulates expression of cell cycle-related genes and causes G_0_/G_1_ arrest in NSCLC^32^; downregulation of METTL7B in ccRCC induces G_0_/G_1_ arrest and apoptosis^45^. Our lipidomic analysis revealed a potential mechanism underlying this cell cycle regulation. Deletion of METTL7B increases ceramide levels and decreases TG levels. Ceramides are well-established tumor suppressors that function by inducing apoptosis and inhibiting cell proliferation through blocking cell cycle transition^58^. The increase in ceramide levels following METTL7B deletion may therefore contribute to the cell cycle arrest and reduced proliferation observed in METTL7B-KO cells. Together, these findings suggest that METTL7B promotes pancreatic cancer progression, at least in part, by modulating lipid metabolism to limit ceramide accumulation and support cell cycle progression, positioning METTL7B as a potential therapeutic target.

In conclusion, our study supports a role for METTL7B as a critical oncogenic factor in pancreatic cancer that is transcriptionally regulated by HNF4A and HNF4G and is functionally linked to LD accumulation, lipid composition, and tumor progression. Given that METTL7B expression is associated with poor patient prognosis, targeting this pathway may represent a promising therapeutic strategy for pancreatic cancer. Future studies are needed to identify the endogenous methylation targets of METTL7B and to elucidate the detailed molecular mechanisms by which METTL7B-mediated lipid metabolism regulates cell cycle progression. These investigations will further advance our understanding of this pathway and its therapeutic potential.

## Materials and Methods

### Cell culture

AsPC-1 cells (ATCC, CRL-1682) were cultured in RPMI-1640 medium (Wako, 189-02025) supplemented with 10% fetal bovine serum (FBS), 1 mM sodium pyruvate (Wako, 190-14881), 10 mM HEPES (Wako, 345-06681), and D(+)-glucose (Wako, 079-05511) at a final concentration of 4.5 g/L. Cells were maintained at 37°C in a humidified atmosphere with 5% CO_2_.

### Expression profiling of methyltransferases in pancreatic cancer

The relative expression levels of 155 methyltransferases in pancreatic cancer tissues compared with normal pancreatic tissues were analyzed using The Cancer Genome Atlas (TCGA)^70^ and the Genotype-Tissue Expression (GTEx) datasets^71^, following the methodology described previously^72^. The list of methyltransferases in humans was obtained from UniProt^73^. Normalized count data (RSEM expected_count normalized using DESeq2) were downloaded from the UCSC Xena platform^74^. The relative expression levels of each methyltransferase in tumor samples were calculated by normalizing the expression level in each sample to the median expression of normal pancreatic tissue samples. The relative expression levels of methyltransferases between tumor and normal tissues were compared using the Wilcoxon test.

### Correlation between methyltransferase expression and patient survival in pancreatic cancer

The association between the expression of methyltransferases and patient survival in pancreatic cancer was analyzed using the TCGA dataset, following the methodology described previously^72^. Patient survival data from the TCGA Pan-Cancer Clinical Data Resource (TCGA-CDR) dataset^75^ were downloaded from UCSC Xena^74^. Patients were categorized into high- and low-expression groups based on the median expression of each methyltransferase. The association between the expression of each methyltransferase and patient survival in pancreatic cancer was evaluated using the Cox proportional hazards model with the coxph function from the survival R package. Kaplan–Meier plots were generated using both the survival and survminer R packages (https://github.com/kassambara/survminer).

### Pan-cancer expression and survival analysis of METTL7B

The relative expression levels of METTL7B in tumor tissues compared with their normal tissues across various cancer types were analyzed using the TCGA^70^ and GTEx datasets^71^. Normalized count data were downloaded from the UCSC Xena platform^74^. The relative expression level of each gene in tumor samples was calculated by normalizing the expression level in each sample to the median expression of normal tissue samples.

The association between the expression of METTL7B and patient survival for each cancer type was analyzed using the TCGA dataset. Patient survival data from the TCGA-CDR dataset^75^ were downloaded from UCSC Xena^74^. Patients were categorized into high- and low-expression groups based on the median of METTL7B. The association between the expression of METTL7B and patient survival in each cancer type was evaluated using the Cox proportional hazards model with the coxph function from the survival R package.

### METTL7B knockout cells

Two guide RNA (gRNA) sequences targeting the region near the METTL7B start codon were designed using Benchling and inserted into the BbsI restriction site of pSpCas9(BB)-2A-GFP (PX458; Addgene, #48138)^76^ to generate two separate plasmids. The plasmids were transfected into AsPC-1 cells using Lipofectamine™ 2000 Transfection Reagent (ThermoFisher Scientific, 11668027). After 24 hours, cells were detached with trypsin-EDTA, and GFP-positive cells were isolated by fluorescence-activated cell sorting (FACS) using a FACSAria™ II cell sorter (BD Biosciences). GFP-positive cells were then seeded into 96-well plates by limiting dilution to obtain single-cell clones. Following expansion, METTL7B knockout clones were identified by western blot analysis using an anti-METTL7B antibody, and clones showing loss of METTL7B protein expression (designated 7B KO2 and 7B KO3) were selected.

### Metastasis Assay

To evaluate tumorigenic potential in vivo, BALB/c nude mice (CAnN.Cg-Foxn1nu/CrlCrlj) were administered an anti-asialo GM1 antibody (Fujifilm Wako, Tokyo, Japan) by intraperitoneal injection 2 hours prior to tumor cell inoculation at a dose of 20 µg/head. Subsequently, 1 × 10⁶ cells of each AsPC-1 clone, suspended in 500 µL of PBS, were injected into the spleen. Immediately after cell injection, splenectomy was performed using an electric cautery device. Following abdominal closure, mice were maintained for three weeks before being sacrificed. After euthanasia, the liver was perfused and fixed with 10% neutral buffered formalin and excised. For each mouse, four liver sections were prepared as formalin-fixed paraffin-embedded (FFPE) specimens and stained with hematoxylin and eosin (H&E). This procedure was performed using six mice per clone. Whole-slide images were generated from the H&E-stained liver sections using a digital slide scanner (NanoZoomer; Hamamatsu Photonics, Hamamatsu, Japan). For each mouse, four representative liver sections were examined, and tumor regions were manually annotated by a pathologist. Tumor burden was quantified as the percentage of tumor area relative to the total liver area using QuPath software^77^.

### Western Blotting

Cells were washed with PBS and lysed for 30 min at 4°C with lysis buffer (50 mM HEPES [pH 7.5], 150 mM KCl, 0.5% NP40, 2 mM EDTA, 1 mM NaF, protease inhibitor cocktail [Sigma-Aldrich, P8340] diluted 1:1000). After centrifugation at 16,100 × g for 30 min at 4°C, the protein concentration of lysates was determined using the Takara BCA Protein Assay Kit (Takara, T9300A). All lysates were boiled for 5 min. The boiled samples were resolved by SDS-PAGE using 4%–15% Mini-PROTEAN TGX Precast Protein Gels (Bio-Rad, #4561086) and transferred to polyvinylidene difluoride (PVDF) membranes (Merck Millipore, Immobilon-P, #IPVH00010). After blocking with Bullet Blocking One for Western Blotting (Nacalai Tesque, #13779-01) for 5 min at room temperature (RT), membranes were incubated with anti-METTL7B antibody (Atlas Antibodies, HPA038644, 1:2000 dilution) or anti-ACTB antibody (MBL, M177-3, 1:5000 dilution) diluted in IMMUNO SHOT Reagent 1 (Cosmo Bio, #IS-001-250) overnight at 4°C. After washing with TBST (20 mM Tris-HCl [pH 7.5], 150 mM NaCl, 0.1% Tween-20), membranes were incubated with HRP-conjugated anti-rabbit IgG antibody (Nacalai Tesque, #21858-24, 1:5000 dilution) or HRP-conjugated anti-mouse IgG antibody (Nacalai Tesque, #21860-74, 1:5000 dilution) diluted in IMMUNO SHOT Reagent 2 (Cosmo Bio, #IS-002-250) for 1 hour at RT. After washing with TBST, signals were visualized using Immobilon Forte Western HRP substrate (Merck Millipore, #WBKLS0500) and the LAS-4000UVmini Luminescent Image Analyzer (Fujifilm).

### Cell proliferation assay

Cell proliferation was assessed using the CellTiter-Glo 2.0 Cell Viability Assay Kit (Promega, G9241) according to the manufacturer’s protocol with minor modifications. Cells were seeded in 96-well plates 24 hours before the assay. At 0, 48, and 96 hours, CellTiter-Glo 2.0 Reagent diluted 1:1 in PBS was added to each well at a volume equal to the culture medium (100 µL). Plates were incubated for 10 min at RT, and luminescence was measured using a GloMax Discover Microplate Reader (Promega, GM3000).

### Wound healing assay

A wound healing assay was performed using 4-well Culture-Inserts (ibidi, ib80469). Inserts were placed on the bottom of each well of a 12-well plate, and cells were seeded into each well of the inserts. After overnight incubation, inserts were removed to create wounds. Wound closure was monitored over time using a microscope (Leica, DMI6000B). Wound area was quantified using the Wound Healing Size Tool ^78^ in Fiji ^79^.

### RNA-seq

Total RNA was isolated using a Nucleospin RNA Clean-up kit (MACHEREY-NAGEL, #740948). To remove genomic DNA contamination, RNA samples were treated with Recombinant DNase I (Takara, #2270A) and Recombinant RNase Inhibitor (Takara, #2313A) for 20 min at 37°C, followed by purification using a Nucleospin RNA Clean-up kit (MACHEREY-NAGEL, #740948). Strand-specific mRNA library preparation and sequencing were performed by BGI Genomics using the DNBSEQ platform to generate paired-end reads.

### RNA-seq analysis

FASTQ files were preprocessed using fastp (version 0.23.2)^80^, and the preprocessed FASTQ files were mapped to the human reference genome (GRCh38.p13) using STAR (version 2.7.10a)^81^ with annotations from GENCODE (Human Release 43). The number of reads mapped to each transcript was counted using StringTie (version 2.2.1)^82^, and the transcript-level counts were summarized to gene-level counts using tximport (version 1.32.0)^83^. Differential gene expression analysis was performed using DESeq2 (version 1.44.0)^84^, and only genes with a mean DESeq2 normalized count of at least 100 were included in downstream analyses. Functional enrichment analysis was performed using the gost function from the gprofiler2 package (version 0.2.3)^85^. Gene Ontology (GO) biological process (BP) terms significantly associated with the differentially expressed genes were clustered using simplifyEnrichment (version 1.14.1)^86^.

### Search for transcription factors regulating METTL7B expression

To identify transcription factors (TFs) regulating METTL7B expression, we used a two-pronged approach. First, TFs that bind near the transcription start site (TSS) of METTL7B were identified using ChIP-Atlas^87^. For each TF, the maximum fold-enrichment value across all ChIP-seq datasets was determined. Second, correlation data between METTL7B expression and each gene in pancreatic cancer patients were obtained from the ‘Pancreatic Adenocarcinoma (TCGA, Firehose Legacy)’ dataset via the cBioPortal^88^. TFs that met both criteria—binding near the METTL7B TSS and showing expression correlation with METTL7B in pancreatic cancer—were identified as candidate regulators. Expression data from the Australian Pancreatic Cancer Genome Initiative (APGI) dataset^89^ were downloaded from Bailey et al. (https://www.nature.com/articles/nature16965#Sec44). For pan-tissue correlation analysis of METTL7B and HNF4A/HNF4G and the subtype analysis of pancreatic cancer samples, the TCGA^90^ and GTEx^71^ datasets were used via the UCSC Xena platform^74^.

### Knockdown and inhibition of HNF4A and HNF4G

Small interfering RNA (siRNA) duplexes targeting HNF4A or HNF4G were purchased from Thermo Fisher Scientific. Cells were transfected with siRNAs using Lipofectamine RNAiMAX Transfection Reagent (Thermo Fisher Scientific, 13778500) according to the manufacturer’s protocol. The siRNA sequences are listed in Supplementary Table S4. Alternatively, HNF4 activity was inhibited by treating AsPC-1 cells with HNF4 inhibitors BIM5078 (Sigma-Aldrich, SML0528) or BI6015 (Cayman, 12032) at 5 µM, or with DMSO as a control. Cells were incubated with inhibitors for 72 hours before RNA isolation.

### qRT-PCR analysis

Total RNA was isolated using the EconoSpin^TM^ for RNA (Ajinomoto Bio-Pharma, EP-21201). Genomic DNA removal and reverse transcription to cDNA were performed using the PrimeScript RT Reagent kit with gDNA eraser (Takara, #RR047B). Quantitative real-time PCR (qRT-PCR) was performed using TB Green Premix Ex Taq II (Takara Bio, #RR820L) on a Thermal Cycler Dice Real-Time System (Takara Bio). Primer sequences are listed in Supplementary Table S5.

### Immunostaining

After washing with PBS, cells were fixed in glyoxal fixative solution^91^ (2.84 ml H_2_O, 789 µl ethanol, 313 µl 40% glyoxal, 30 µl acetic acid) at room temperature for 20 min. After washing three times with PBS containing 0.1% Tween-20 (PBS-T), cells were permeabilized by treatment with PBS containing 0.1% Triton X-100 for 5 minutes at 4°C. After washing three times with PBS-T, samples were blocked with PBS containing 1% BSA and 0.1% Tween-20 (BSA-T) for 1 hour. Samples were then incubated at 4°C overnight with primary antibodies against METTL7B (Atlas Antibodies, HPA038644, 1:200 dilution), PDIA3 (Sigma-Aldrich, AMAB90988, 1:200 dilution), or CD29 (BD, 610467, 1:200 dilution) diluted in BSA-T. After washing three times with PBS-T, samples were incubated at room temperature for 1 hour with Alexa Fluor 488-labeled anti-rabbit IgG secondary antibody (Thermo Fisher Scientific, A21206, 1:1000 dilution), Alexa Fluor 555-labeled anti-mouse IgG secondary antibody (Thermo Fisher Scientific, A31570, 1:1000 dilution), or Alexa Fluor 647-labeled anti-mouse IgG secondary antibody (Thermo Fisher Scientific, A31571, 1:1000 dilution) diluted in BSA-T, followed by three washes with PBS-T. For LD staining, samples were incubated with 0.1 µM Lipi-Blue (Dojindo, LD01) or Lipi-Deep Red (Dojindo, LD04) in PBS at 37°C for 1 hour. For nuclear staining, samples were incubated with 1 µg/mL DAPI solution (Dojindo, D523) in PBS for 10 minutes at room temperature. After washing three times with PBS, samples were mounted with ProLong™ Gold Antifade Mountant (Thermo Fisher Scientific, P36934) for imaging. Images were acquired with a 63× oil immersion objective in AiryScan mode on an LSM900 microscope (Zeiss). AiryScan processing was performed on the acquired images.

### Quantification of LDs

LDs were quantified using Fiji^79^. First, maximum intensity Z-projection was performed on the LD channel. The Z-projected images were then binarized using the Yen algorithm. Noise was removed by performing three iterations of erosion and three iterations of dilation processes on the binarized images. Watershed segmentation was then applied to the denoised images to separate contacting LDs. Particle analysis was performed on the watershed-processed images to measure the size of each LD. Finally, the number of LDs per cell was quantified based on cell regions defined using the CD29 channel.

### Lipidome/Metabolome

Lipidomic and metabolomic profiling was performed by liquid chromatography–mass spectrometry (LC-MS) following extraction of cell pellets with acetonitrile, as previously described for hydrophilic interaction liquid chromatography (HILIC)-based metabolomics^92^ and C18-based lipidomics^93^. Approximately 5 × 10^6^ cells were scraped and pelleted. After adding 100 µl of acetonitrile, samples were mixed for 30 sec and homogenized for 5 min at 4°C. After centrifugation at 16,400 × g for 10 min at 4°C, the supernatant was transferred to the sample vial for LC/MS. An aliquot (20 µl) from each sample was mixed for quality control (QC). The supernatants were analyzed using ultra-high-performance liquid chromatography–Fourier transform mass spectrometry (UHPLC-FTMS) in four modes: metabolomics using a HILIC column (positive and negative ion modes) and lipidomics using a C18 column (positive and negative ion modes). Pre-processing including peak picking and quantification was performed using ProgenesisQI (Waters).

All downstream analyses were performed using custom R scripts. Quality control filtering was performed by calculating median peak intensities for each peak across replicates within each cell type and analytical mode. Peaks were retained only if their QC sample median values fell within the range defined by the minimum and maximum median values across all cell types for that peak and analytical mode. Peak intensity changes for each knockout condition were classified as upregulated, downregulated, or unchanged based on statistical significance (*p* < 0.05) and the direction of log_2_ fold change. Peaks were further classified based on common change patterns between the two independent METTL7B-KO clones as commonly upregulated, commonly downregulated, or other combinations.

Lipid class estimation was performed using annotations from the LIPID MAPS Structure Database (LMSD)^94,95^ assigned to each peak based on mass-to-charge ratio (m/z) matching. For peaks with multiple LMSD ID matches, the fraction of each main class (MAIN_CLASS) and subclass (SUB_CLASS) category was calculated as the proportion of matching IDs belonging to that category relative to the total number of matches for each peak and analytical mode. Lipid composition was estimated by calculating the percentage of each main class and subclass category for all peaks, upregulated peaks, and downregulated peaks, both within each analytical mode and across all analytical modes combined.

For chemical formula analysis, ceramide, TG, and DG peaks with main class fractions > 0.5 and a single chemical formula annotation were used. For TG-specific analysis, an additional filter was applied: only peaks detected in lipidomics positive mode with retention times > 10 min were included to exclude peaks that were incorrectly annotated as TGs. For analysis across lipid classes, retention time filtering was not performed, even for TGs. From the chemical formulas, degrees of unsaturation and total carbon numbers were calculated. The degree of unsaturation was determined using the equation: (2C + 2 – H + N – halogens) / 2.

## Data availability

RNA-seq data has been deposited to the DNA Data Bank of Japan (DDBJ) Sequence Read Archive (DRA; https://www.ddbj.nig.ac.jp/dra/) under accession numbers DRR932710-DRR932715. Source data are provided with this paper.

## Competing interest statement

The authors declare that they have no competing interests.

## Acknowledgements

We thank Y. Uchijima for technical assistance with FACS. This work was supported by the Japan Society for the Promotion of Science (JSPS) [Grant Number JP21H02758 (K.Taniue), JP21K19402 (K.Taniue), JP21H04792 (N.A.), JP22KK0285 (K.Taniue), JP22KK0125 (Y.M), JP22KJ0965 (S.M.), JP24KJ2206 (S.M.), JP 24K02431 (Y.M), and JP24K22066 (K.Taniue)]. N.A. and K.Taniue were supported by the Takeda Science Foundation and the Uehara Memorial Foundation. Y.M. and K.Taniue were supported by the Naito Foundation. Y.M was supported by the Princess Takamatsu Cancer Research Fund and the Project Mirai Cancer Research Grants. K.Taniue was supported by the Japan Foundation for Applied Enzymology, the ONO Medical Research Foundation, the Astellas Foundation for Research on Metabolic Disorders, the MSD Life Science Foundation, and the Kobayashi Foundation. Part of the computations were performed on the NIG supercomputer at ROIS National Institute of Genetics. The results shown here are in part based upon data generated by the TCGA Research Network: https://www.cancer.gov/tcga.

## Author Contributions

S.M., K.Taniue, and N.A. designed the research. S.M. and K.Taniue performed experiments, except lipidomic and metabolomic analyses, with assistance from A.N., O. K.Takahashi and Y.M. D.S. and J.A. performed lipidomic and metabolomic experiments. N.T., H.T., A.S., K.Takahashi and Y.M. performed metastasis assay. S.M. performed bioinformatic analyses. S.M., K.Taniue, and N.A. wrote and revised the manuscript.

## Notes

### Competing Interest Statement

The authors have declared no competing interest.

## References

1. Hanahan, D. & Weinberg, R. A. Hallmarks of Cancer: The Next Generation. Cell 144, 646–674 (2011).

2. Snaebjornsson, M. T., Janaki-Raman, S. & Schulze, A. Greasing the Wheels of the Cancer Machine: The Role of Lipid Metabolism in Cancer. Cell Metabolism vol. 31 Preprint at 10.1016/j.cmet.2019.11.010 (2020).

3. Broadfield, L. A., Pane, A. A., Talebi, A., Swinnen, J. V. & Fendt, S. M. Lipid metabolism in cancer: New perspectives and emerging mechanisms. Developmental Cell vol. 56 Preprint at 10.1016/j.devcel.2021.04.013 (2021).

4. Petrov, M. S. & Taylor, R. Intra-pancreatic fat deposition: bringing hidden fat to the fore. Nature Reviews Gastroenterology and Hepatology vol. 19 Preprint at 10.1038/s41575-021-00551-0 (2022).

5. Wagner, R. et al. Metabolic implications of pancreatic fat accumulation. Nature Reviews Endocrinology vol. 18 Preprint at 10.1038/s41574-021-00573-3 (2022).

6. Sreedhar, U. L., DeSouza, S. V., Park, B. & Petrov, M. S. A Systematic Review of Intra-pancreatic Fat Deposition and Pancreatic Carcinogenesis. Journal of Gastrointestinal Surgery 24, (2020).

7. Desai, V. et al. Pancreatic Fat Infiltration Is Associated with a Higher Risk of Pancreatic Ductal Adenocarcinoma. Visc. Med. 36, (2020).

8. Hori, M. et al. Enhancement of carcinogenesis and fatty infiltration in the pancreas in n-nitrosobis(2-oxopropyl)amine-treated hamsters by high-fat diet. in Pancreas vol. 40 (2011).

9. Philip, B. et al. A high-fat diet activates oncogenic Kras and COX2 to induce development of pancreatic ductal adenocarcinoma in mice. Gastroenterology 145, (2013).

10. Yamazaki, H. et al. Evidence for a causal link between intra-pancreatic fat deposition and pancreatic cancer: A prospective cohort and Mendelian randomization study. Cell Rep. Med. 5, (2024).

11. Olzmann, J. A. & Carvalho, P. Dynamics and functions of lipid droplets. Nature Reviews Molecular Cell Biology vol. 20 Preprint at 10.1038/s41580-018-0085-z (2019).

12. Thiam, A. R., Farese, R. V. & Walther, T. C. The biophysics and cell biology of lipid droplets. Nature Reviews Molecular Cell Biology vol. 14 Preprint at 10.1038/nrm3699 (2013).

13. Mathiowetz, A. J. & Olzmann, J. A. Lipid droplets and cellular lipid flux. Nature Cell Biology vol. 26 Preprint at 10.1038/s41556-024-01364-4 (2024).

14. Cruz, A. L. S., Barreto, E. de A., Fazolini, N. P. B., Viola, J. P. B. & Bozza, P. T. Lipid droplets: platforms with multiple functions in cancer hallmarks. Cell Death and Disease vol. 11 Preprint at 10.1038/s41419-020-2297-3 (2020).

15. Greenberg, A. S. et al. The role of lipid droplets in metabolic disease in rodents and humans. Journal of Clinical Investigation vol. 121 Preprint at 10.1172/JCI46069 (2011).

16. Rozeveld, C. N., Johnson, K. M., Zhang, L. & Razidlo, G. L. KRAS controls pancreatic cancer cell lipid metabolism and invasive potential through the lipase HSL. Cancer Res. 80, (2020).

17. Sunami, Y., Rebelo, A. & Kleeff, J. Lipid metabolism and lipid droplets in pancreatic cancer and stellate cells. Cancers vol. 10 Preprint at 10.3390/cancers10010003 (2018).

18. Michalak, E. M., Burr, M. L., Bannister, A. J. & Dawson, M. A. The roles of DNA, RNA and histone methylation in ageing and cancer. Nature Reviews Molecular Cell Biology vol. 20 Preprint at 10.1038/s41580-019-0143-1 (2019).

19. Herz, H. M., Garruss, A. & Shilatifard, A. SET for life: Biochemical activities and biological functions of SET domain-containing proteins. Trends in Biochemical Sciences vol. 38 Preprint at 10.1016/j.tibs.2013.09.004 (2013).

20. Falnes, P. Closing in on human methylation—the versatile family of seven-β-strand (METTL) methyltransferases. Nucleic Acids Res. 52, (2024).

21. Greenberg, M. V. C. & Bourc’his, D. The diverse roles of DNA methylation in mammalian development and disease. Nature Reviews Molecular Cell Biology vol. 20 Preprint at 10.1038/s41580-019-0159-6 (2019).

22. Michalak, E. M., Burr, M. L., Bannister, A. J. & Dawson, M. A. The roles of DNA, RNA and histone methylation in ageing and cancer. Nature Reviews Molecular Cell Biology vol. 20 Preprint at 10.1038/s41580-019-0143-1 (2019).

23. Li, G., et al. Critical roles and clinical perspectives of RNA methylation in cancer. MedComm vol. 5 Preprint at 10.1002/mco2.559 (2024).

24. Hamamoto, R. & Nakamura, Y. Dysregulation of protein methyltransferases in human cancer: An emerging target class for anticancer therapy. Cancer Science vol. 107 Preprint at 10.1111/cas.12884 (2016).

25. Sanderson, S. M., Gao, X., Dai, Z. & Locasale, J. W. Methionine metabolism in health and cancer: a nexus of diet and precision medicine. Nature Reviews Cancer vol. 19 Preprint at 10.1038/s41568-019-0187-8 (2019).

26. Maldonato, B. J., Russell, D. A. & Totah, R. A. Human METTL7B is an alkyl thiol methyltransferase that metabolizes hydrogen sulfide and captopril. Sci. Rep. 11, (2021).

27. Russell, D. A. et al. METTL7A (TMT1A) and METTL7B (TMT1B) Are Responsible for Alkyl S-Thiol Methyl Transferase Activity in Liver. Drug Metabolism and Disposition 51, (2023).

28. Robey, R. W. et al. The Methyltransferases METTL7A and METTL7B Confer Resistance to Thiol-Based Histone Deacetylase Inhibitors. Mol. Cancer Ther. 23, (2024).

29. McKinnon, C. M. & Mellor, H. The tumor suppressor RhoBTB1 controls Golgi integrity and breast cancer cell invasion through METTL7B. BMC Cancer 10.1186/s12885-017-3138-3 (2017) doi:10.1186/s12885-017-3138-3.

30. Ye, D. et al. METTL7B promotes migration and invasion in thyroid cancer through epithelial-mesenchymal transition. J. Mol. Endocrinol. 10.1530/JME-18-0261 (2019) doi:10.1530/JME-18-0261.

31. Zhu, J. et al. circ-PSD3 promoted proliferation and invasion of papillary thyroid cancer cells via regulating the miR-7-5p/METTL7B axis. Journal of Receptors and Signal Transduction 42, (2022).

32. Liu, D. et al. METTL7B Is Required for Cancer Cell Proliferation and Tumorigenesis in Non-Small Cell Lung Cancer. Front. Pharmacol. 11, (2020).

33. Li, W. et al. Downregulation of METTL7B Inhibits Proliferation of Human Clear Cell Renal Cancer Cells In Vivo and In Vitro. Front. Oncol. 11, (2021).

34. Ali, J. et al. METTL7B (methyltransferase-like 7B) identification as a novel biomarker for lung adenocarcinoma. Ann. Transl. Med. 10.21037/atm-20-4574 (2020) doi:10.21037/atm-20-4574.

35. Li, R., Mu, C., Cao, Y. & Fan, Y. METTL7B serves as a prognostic biomarker and promotes metastasis of lung adenocarcinoma cells. Ann. Transl. Med. 10, (2022).

36. Fu, R., Luo, X., Ding, Y. & Guo, S. Prognostic Potential of METTL7B in Glioma. Neuroimmunomodulation 29, (2022).

37. Jiang, Z. et al. METTL7B is a novel prognostic biomarker of lower-grade glioma based on pan-cancer analysis. Cancer Cell Int. 21, (2021).

38. Chen, X. et al. Characterization of METTL7B to Evaluate TME and Predict Prognosis by Integrative Analysis of Multi-Omics Data in Glioma. Front. Mol. Biosci. 8, (2021).

39. Turró, S. et al. Identification and characterization of associated with lipid droplet protein 1: A novel membrane-associated protein that resides on hepatic lipid droplets. Traffic 7, (2006).

40. Franjic, D. et al. Transcriptomic taxonomy and neurogenic trajectories of adult human, macaque, and pig hippocampal and entorhinal cells. Neuron 110, (2022).

41. Bersuker, K. et al. A Proximity Labeling Strategy Provides Insights into the Composition and Dynamics of Lipid Droplet Proteomes. Dev. Cell 44, (2018).

42. Halbrook, C. J., Lyssiotis, C. A., Pasca di Magliano, M. & Maitra, A. Pancreatic cancer: Advances and challenges. Cell vol. 186 Preprint at 10.1016/j.cell.2023.02.014 (2023).

43. Xu, L. et al. METTL7B contributes to the malignant progression of glioblastoma by inhibiting EGR1 expression. Metab. Brain Dis. 37, (2022).

44. Xiong, Y., Li, M., Bai, J., Sheng, Y. & Zhang, Y. High Level of METTL7B Indicates Poor Prognosis of Patients and Is Related to Immunity in Glioma. Front. Oncol. 11, (2021).

45. Li, W. et al. Downregulation of METTL7B Inhibits Proliferation of Human Clear Cell Renal Cancer Cells In Vivo and In Vitro. Front. Oncol. 11, (2021).

46. Lücke, J. et al. Protocol for generating lung and liver metastasis in mice using models that bypass intravasation. STAR Protoc. 5, (2024).

47. O’Brien, M., Ernst, M. & Poh, A. R. An intrasplenic injection model of pancreatic cancer metastasis to the liver in mice. STAR Protoc. 4, (2023).

48. Sladek, F. M., Zhong, W., Lai, E. & Darnell, J. E. Liver-enriched transcription factor HNF-4 is a novel member of the steroid hormone receptor superfamily. Genes Dev. 4, (1990).

49. Palanker, L., Tennessen, J. M., Lam, G. & Thummel, C. S. Drosophila HNF4 Regulates Lipid Mobilization and β-Oxidation. Cell Metab. 9, (2009).

50. Storelli, G., Nam, H. J., Simcox, J., Villanueva, C. J. & Thummel, C. S. Drosophila HNF4 Directs a Switch in Lipid Metabolism that Supports the Transition to Adulthood. Dev. Cell 48, (2019).

51. Chen, L. et al. HNF4 Regulates Fatty Acid Oxidation and Is Required for Renewal of Intestinal Stem Cells in Mice. Gastroenterology 158, (2020).

52. Hayhurst, G. P., Lee, Y.-H., Lambert, G., Ward, J. M. & Gonzalez, F. J. Hepatocyte Nuclear Factor 4α (Nuclear Receptor 2A1) Is Essential for Maintenance of Hepatic Gene Expression and Lipid Homeostasis. Mol. Cell. Biol. 21, (2001).

53. Kiselyuk, A. et al. HNF4α antagonists discovered by a high-throughput screen for modulators of the human insulin promoter. Chem. Biol. 19, (2012).

54. Wu, C. C. et al. Organellar proteomics reveals Golgi arginine dimethylation. Mol. Biol. Cell 15, (2004).

55. Ayari, S. et al. Hnf4g invalidation prevents diet-induced obesity via intestinal lipid malabsorption. J. Endocrinol. 252, (2021).

56. Vonolfen, M. C. et al. Drosophila HNF4 acts in distinct tissues to direct a switch between lipid storage and export in the gut. Cell Rep. 43, (2024).

57. Girard, R. et al. The transcription factor hepatocyte nuclear factor 4A acts in the intestine to promote white adipose tissue energy storage. Nat. Commun. 13, (2022).

58. Morad, S. A. F. & Cabot, M. C. Ceramide-orchestrated signalling in cancer cells. Nature Reviews Cancer vol. 13 Preprint at 10.1038/nrc3398 (2013).

59. Ogretmen, B. Sphingolipid metabolism in cancer signalling and therapy. Nature Reviews Cancer vol. 18 Preprint at 10.1038/nrc.2017.96 (2017).

60. Ho, Q. W. C., Zheng, X. & Ali, Y. Ceramide Acyl Chain Length and Its Relevance to Intracellular Lipid Regulation. International Journal of Molecular Sciences vol. 23 Preprint at 10.3390/ijms23179697 (2022).

61. Tauchi-Sato, K., Ozeki, S., Houjou, T., Taguchi, R. & Fujimoto, T. The surface of lipid droplets is a phospholipid monolayer with a unique fatty acid composition. Journal of Biological Chemistry 277, (2002).

62. Moessinger, C., Kuerschner, L., Spandl, J., Shevchenko, A. & Thiele, C. Human lysophosphatidylcholine acyltransferases 1 and 2 are located in lipid droplets where they catalyze the formation of phosphatidylcholine. Journal of Biological Chemistry 286, (2011).

63. Wölk, M. & Fedorova, M. The lipid droplet lipidome. FEBS Letters vol. 598 Preprint at 10.1002/1873-3468.14874 (2024).

64. Petrov, M. S. Fatty change of the pancreas: the Pandora’s box of pancreatology. The Lancet Gastroenterology and Hepatology vol. 8 Preprint at 10.1016/S2468-1253(23)00064-X (2023).

65. Tauchi-Sato, K., Ozeki, S., Houjou, T., Taguchi, R. & Fujimoto, T. The surface of lipid droplets is a phospholipid monolayer with a unique fatty acid composition. Journal of Biological Chemistry 277, (2002).

66. Chen, L. et al. A reinforcing HNF4–SMAD4 feed-forward module stabilizes enterocyte identity. Nature Genetics vol. 51 Preprint at 10.1038/s41588-019-0384-0 (2019).

67. Lei, J., Li, G., Yu, H. & An, T. Potent necrosis effect of methanethiol mediated by METTL7B enzyme bioactivation mechanism in 16HBE cell. Ecotoxicol. Environ. Saf. 236, 113486 (2022).

68. Giustarini, D., Dalle-Donne, I., Milzani, A., Fanti, P. & Rossi, R. Analysis of GSH and GSSG after derivatization with N-ethylmaleimide. Nat. Protoc. 8, (2013).

69. Li, X., Gluth, A., Zhang, T. & Qian, W. J. Thiol redox proteomics: Characterization of thiol-based post-translational modifications. Proteomics vol. 23 Preprint at 10.1002/pmic.202200194 (2023).

70. Weinstein, J. N. et al. The cancer genome atlas pan-cancer analysis project. Nat. Genet. 45, (2013).

71. Lonsdale, J., et al. The Genotype-Tissue Expression (GTEx) project. Nature Genetics vol. 45 Preprint at 10.1038/ng.2653 (2013).

72. Mitsutomi, S. et al. Nanopore direct RNA sequencing reveals METTL2A-mediated m3C sites in poly(A) RNA. Genome Res. 35, (2025).

73. Bateman, A. et al. UniProt: the Universal Protein Knowledgebase in 2023. Nucleic Acids Res. 51, (2023).

74. Goldman, M. J. et al. Visualizing and interpreting cancer genomics data via the Xena platform. Nature Biotechnology vol. 38 Preprint at 10.1038/s41587-020-0546-8 (2020).

75. Liu, J. et al. An Integrated TCGA Pan-Cancer Clinical Data Resource to Drive High-Quality Survival Outcome Analytics. Cell 173, (2018).

76. Ran, F. A. et al. Genome engineering using the CRISPR-Cas9 system. Nature Protocols 2013 8:11 8, 2281–2308 (2013).

77. Bankhead, P. et al. QuPath: Open source software for digital pathology image analysis. Sci. Rep. 7, (2017).

78. Suarez-Arnedo, A. et al. An image J plugin for the high throughput image analysis of in vitro scratch wound healing assays. PLoS One 15, (2020).

79. Schindelin, J., et al. Fiji: An open-source platform for biological-image analysis. Nature Methods vol. 9 Preprint at 10.1038/nmeth.2019 (2012).

80. Chen, S., Zhou, Y., Chen, Y. & Gu, J. Fastp: An ultra-fast all-in-one FASTQ preprocessor. In Bioinformatics vol. 34 (2018).

81. Dobin, A. et al. STAR: Ultrafast universal RNA-seq aligner. Bioinformatics 10.1093/bioinformatics/bts635 (2013) doi:10.1093/bioinformatics/bts635.

82. Pertea, M., Kim, D., Pertea, G. M., Leek, J. T. & Salzberg, S. L. Transcript-level expression analysis of RNA-seq experiments with HISAT, StringTie and Ballgown. Nat. Protoc. 11, (2016).

83. Soneson, C., Love, M. I. & Robinson, M. D. Differential analyses for RNA-seq: transcript-level estimates improve gene-level inferences. F1000Res. 4, (2015).

84. Love, M. I., Huber, W. & Anders, S. Moderated estimation of fold change and dispersion for RNA-seq data with DESeq2. Genome Biol. 15, (2014).

85. Peterson, H., Kolberg, L., Raudvere, U., Kuzmin, I. & Vilo, J. gprofiler2 -- an R package for gene list functional enrichment analysis and namespace conversion toolset g: Profiler. F1000Res. 9, (2020).

86. Gu, Z. & Hübschmann, D. simplifyEnrichment: A Bioconductor Package for Clustering and Visualizing Functional Enrichment Results. Genomics Proteomics Bioinformatics 21, (2023).

87. Oki, S. et al. ChIP-Atlas: a data-mining suite powered by full integration of public ChIP-seq data. EMBO Rep. 19, (2018).

88. Cerami, E. et al. The cBio Cancer Genomics Portal: An open platform for exploring multidimensional cancer genomics data. Cancer Discov. 2, (2012).

89. Bailey, P. et al. Genomic analyses identify molecular subtypes of pancreatic cancer. Nature 531, (2016).

90. TCGA Research Network. The Cancer Genome Atlas Program (TCGA). Https://Www.Cancer.Gov/Tcga (2023).

91. Richter, K. N. et al. Glyoxal as an alternative fixative to formaldehyde in immunostaining and super-resolution microscopy. EMBO J. 37, (2018).

92. Saigusa, D. et al. Establishment of protocols for global metabolomics by LC-MS for biomarker discovery. PLoS One 11, (2016).

93. Saigusa, D. et al. Comparison of kit-based metabolomics with other methodologies in a large cohort, towards establishing reference values. Metabolites 11, (2021).

94. Sud, M. et al. LMSD: LIPID MAPS structure database. Nucleic Acids Res. 35, (2007).

95. Fahy, E. et al. Update of the LIPID MAPS comprehensive classification system for lipids. Journal of Lipid Research vol. 50 Preprint at 10.1194/jlr.R800095-JLR200 (2009).

