## Supplementary Figures for "Thiol methyltransferase METTL7B coordinates lipid metabolism and promotes tumor progression in pancreatic cancer"

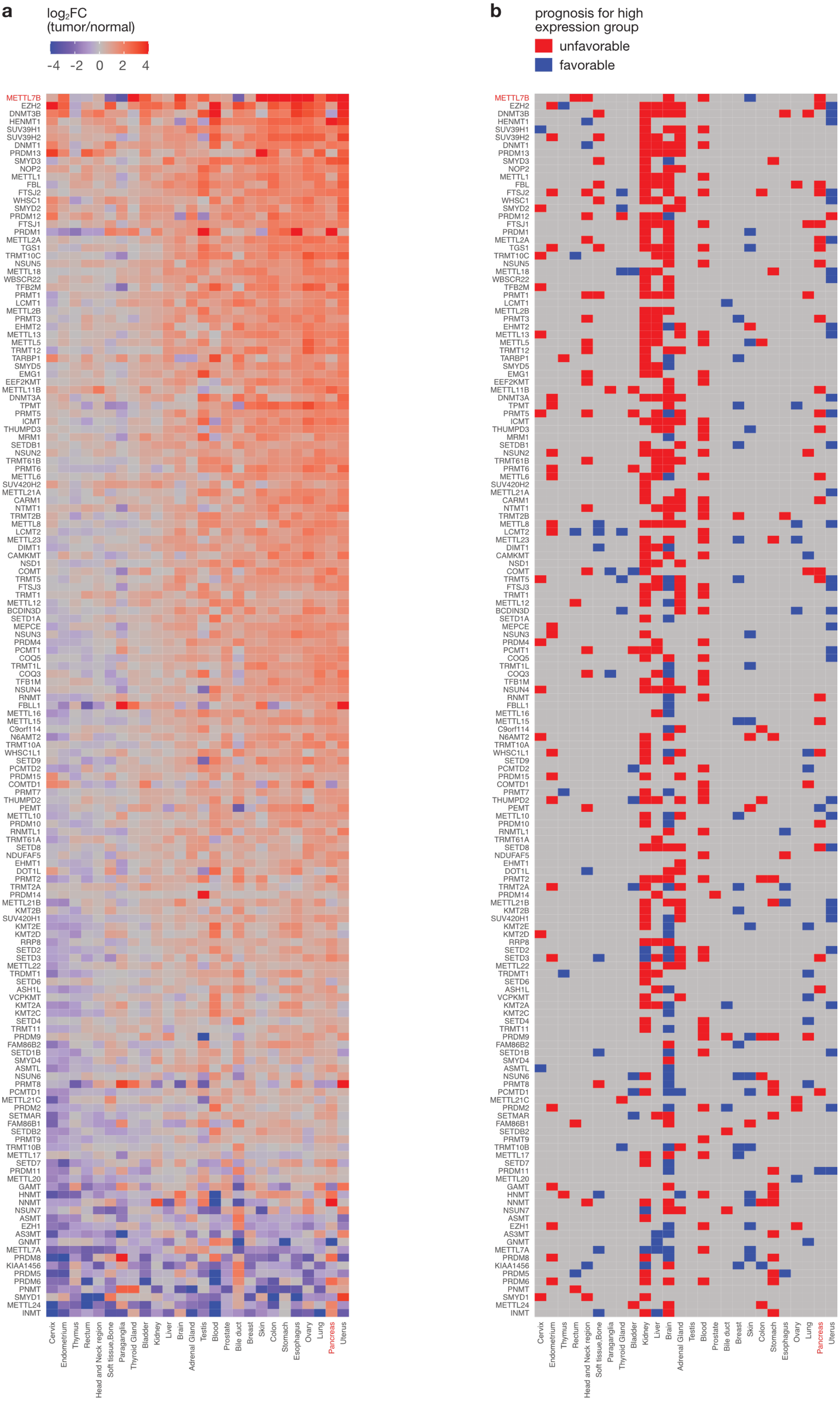
a
log2FC
(tumor/normal)
-4 -2 0 2 4
b
prognosis for high expression group
unfavorable
favorable
METTL7B EZH2 DNMT3B HENMT1 SUV39H1 SUV39H2 DNMT1 PRDM13 SMYD3 NOP2 METTL1 FBL FTSJ2 WHSC1 SMYD2 PRDM12 FTSJ1 PRDM1 METTL2A TGS1 TRMT10C NSUN5 METTL18 WBSCR22 TFB2M PRMT1 LCMT1 METTL2B PRMT3 EHMT2 METTL13 METTL5 TRMT12 TARBP1 SMYD5 EMG1 EEF2KMT METTL11B DNMT3A TPMT PRMT5 ICMT THUMPD3 MRM1 SETDB1 NSUN2 TRMT61B PRMT6 METTL6 SUV420H2 METTL21A CARM1 NTMT1 TRMT2B METTL8 LCMT2 METTL23 DIMT1 CAMKMT NSD1 COMT TRMT5 FTSJ3 TRMT1 METTL12 BCDIN3D SETD1A MEPCE NSUN3 PRDM4 PCMT1 COQ5 TRMT1L COQ3 TFB1M NSUN4 RNMT FBLL1 METTL16 METTL15 C9orf114 N6AMT2 TRMT10A WHSC1L1 SETD9 PCMTD2 PRDM15 COMTD1 PRMT7 THUMPD2 PEMT METTL10 PRDM10 RNMTL1 TRMT61A SETD8 NDUFAF5 EHMT1 DOT1L PRMT2 TRMT2A PRDM14 METTL21B KMT2B SUV420H1 KMT2E KMT2D RRP8 SETD2 SETD3 METTL22 TRDMT1 SETD6 ASH1L VCPKMT KMT2A KMT2C SETD4 TRMT11 PRDM9 FAM86B2 SETD1B SMYD4 ASMTL NSUN6 PRMT8 PCMTD1 METTL21C PRDM2 SETMAR FAM86B1 SETDB2 PRMT9 TRMT10B METTL17 SETD7 PRDM11 METTL20 GAMT HNMT NNMT NSUN7 ASMT EZH1 AS3MT GNMT METTL7A PRDM8 KIAA1456 PRDM5 PRDM6 PNMT SMYD1 METTL24 INMT
Cervix Endometrium Thymus Rectum Head and Neck region Soft tissue,Bone Paraganglia Thyroid Gland Bladder Kidney Liver Brain Adrenal Gland Testis Blood Prostate Bile duct Breast Skin Colon Stomach Esophagus Ovary Lung Pancreas Uterus

**Figure S1 | Pan-cancer landscape of methyltransferase expression and prognosis.**

**a** Heatmap showing $\log_2$ fold change (tumor vs. normal) for 155 methyltransferases across multiple cancer types. *METTL7B* and pancreatic cancer are highlighted in red. Methyltransferases and cancer types are sorted by relative expression in tumor versus normal tissue. **b** Heatmap showing the prognostic association of high methyltransferase with overall survival across multiple cancer types. Red indicates unfavorable prognosis, blue indicates favorable prognosis, and gray indicates no significant association. *METTL7B* and pancreatic cancer are highlighted in red.

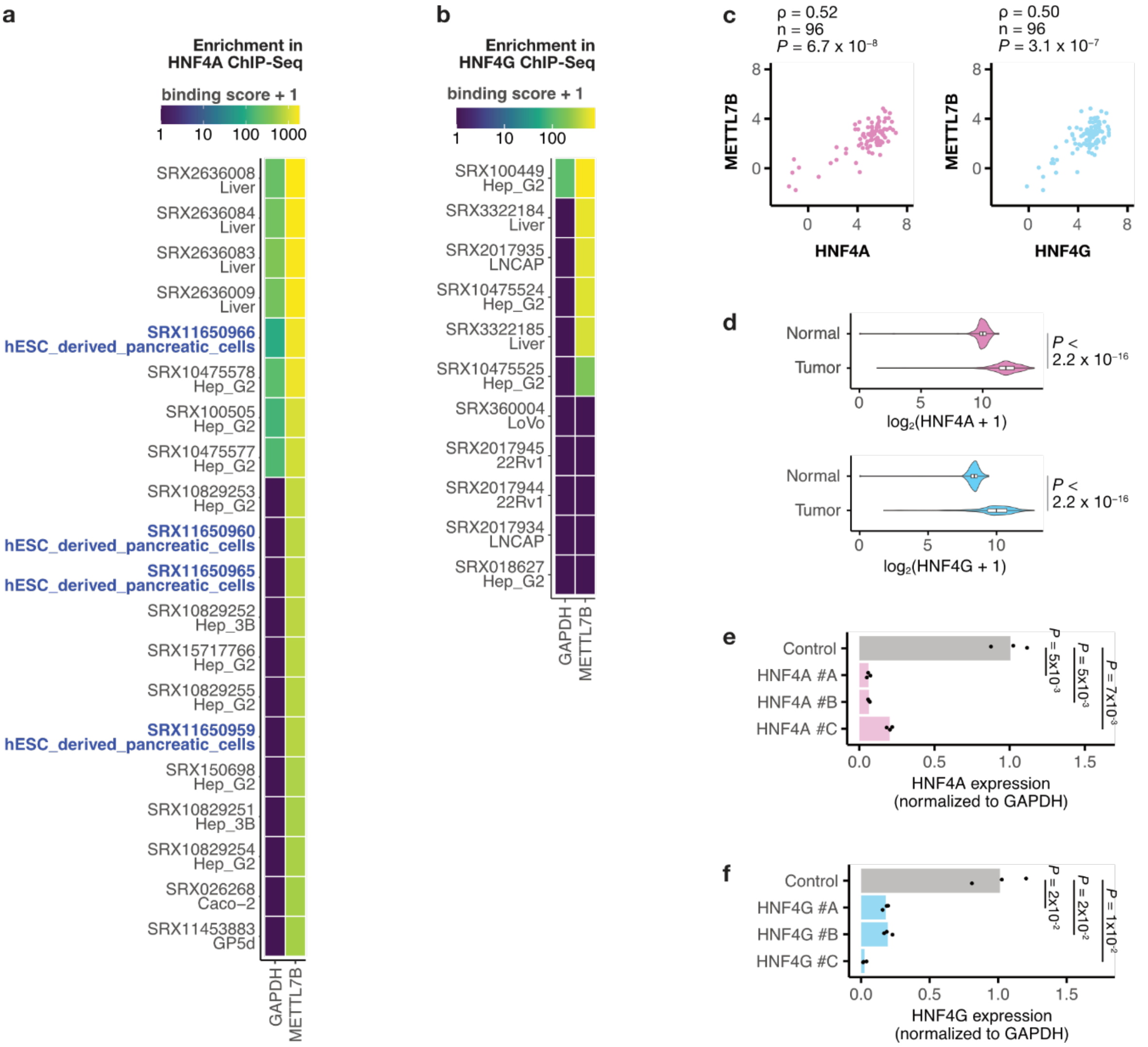


**Figure S2 | HNF4A and HNF4G bind the *METTL7B* locus and regulate its expression**

**a** Heatmap showing HNF4A ChIP-seq binding scores at the *METTL7B* locus and at *GAPDH* (used as a control) across multiple cell lines and tissues, based on ChIP-Atlas data. Pancreatic cell samples are indicated in blue. **b** Heatmap showing HNF4G ChIP-seq binding scores at the *METTL7B* locus and at *GAPDH* (used as a control) across multiple cell lines and tissues, based on ChIP-Atlas data. **c** Scatter plots showing correlations between *METTL7B* expression and *HNF4A* (left) or *HNF4G* (right) expression in the Australian Pancreatic Cancer Genome Initiative (APGI) cohort. n = 96. Spearman's correlation test. **d** Violin plots comparing *HNF4A* (top) and *HNF4G* (bottom) expression levels between normal pancreas (n = 171) and pancreatic tumor (n = 179) samples. Wilcoxon test. **e** Relative *HNF4A* mRNA expression in AsPC-1 cells 72 h after transfection with control or *HNF4A*-targeting siRNAs, normalized to *GAPDH*. Student's *t*-test. n = 3. **f** Relative *HNF4G* mRNA expression in AsPC-1 cells 72 h after transfection with control or *HNF4G*-targeting siRNAs, normalized to *GAPDH*. Student's *t*-test. n = 3.

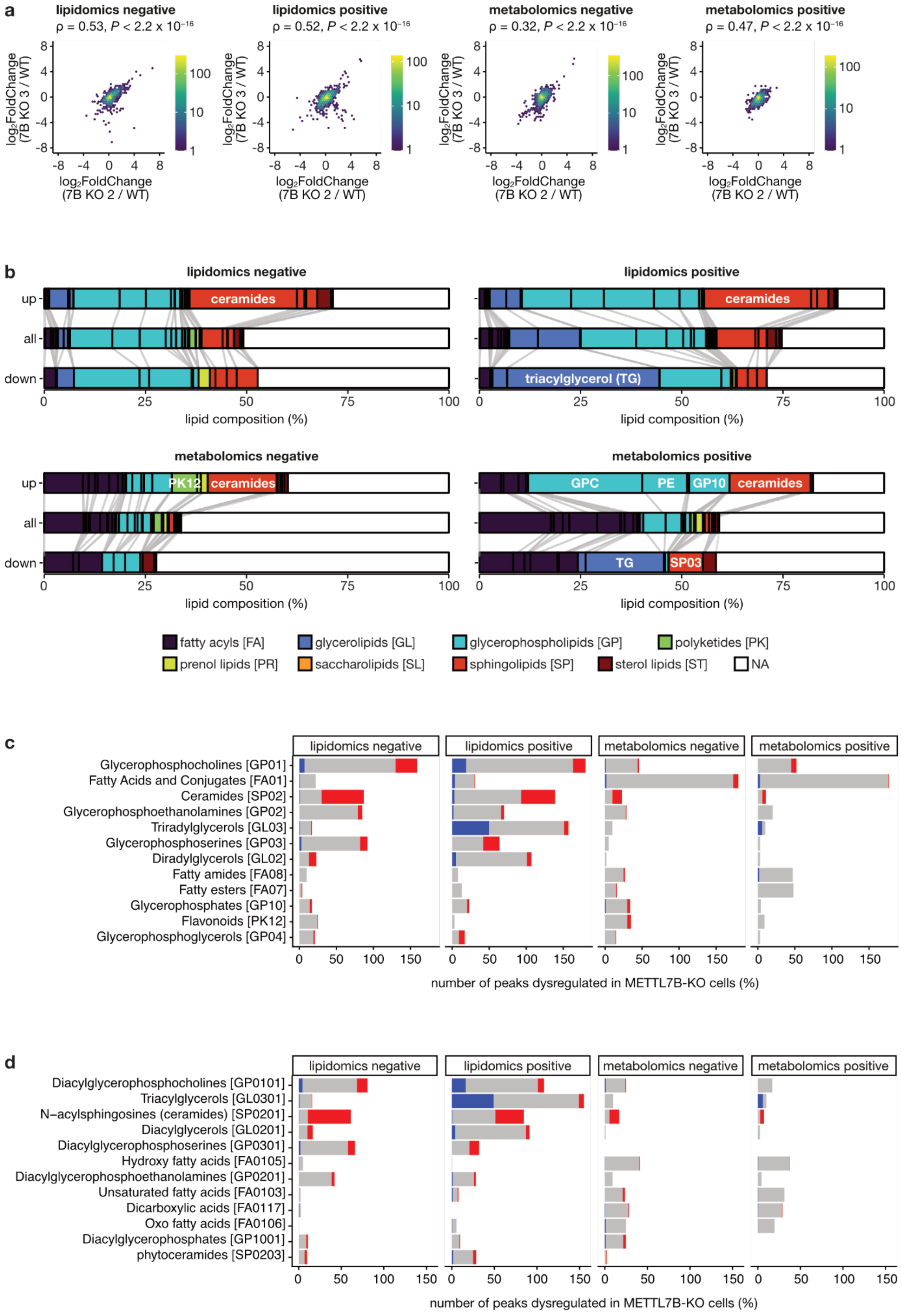
a
lipidomics negative
ρ = 0.53, P < 2.2 x 10⁻¹⁶
lipidomics positive
ρ = 0.52, P < 2.2 x 10⁻¹⁶
metabolomics negative
ρ = 0.32, P < 2.2 x 10⁻¹⁶
metabolomics positive
ρ = 0.47, P < 2.2 x 10⁻¹⁶
log2FoldChange (7B KO 3 / WT)
log2FoldChange (7B KO 2 / WT)
b
lipidomics negative
lipidomics positive
metabolomics negative
metabolomics positive
up
all
down
ceramides
triacylglycerol (TG)
PK12
GPC
PE
GP10
TG
SP03
lipid composition (%)
fatty acyls [FA]
glycerolipids [GL]
glycerophospholipids [GP]
polyketides [PK]
prenol lipids [PR]
saccharolipids [SL]
sphingolipids [SP]
sterol lipids [ST]
NA
c
Glycerophosphocholines [GP01]
Fatty Acids and Conjugates [FA01]
Ceramides [SP02]
Glycerophosphoethanolamines [GP02]
Triradylglycerols [GL03]
Glycerophosphoserines [GP03]
Diradylglycerols [GL02]
Fatty amides [FA08]
Fatty esters [FA07]
Glycerophosphates [GP10]
Flavonoids [PK12]
Glycerophosphoglycerols [GP04]
number of peaks dysregulated in METTL7B-KO cells (%)
d
Diacylglycerophosphocholines [GP0101]
Triacylglycerols [GL0301]
N−acylsphingosines (ceramides) [SP0201]
Diacylglycerols [GL0201]
Diacylglycerophosphoserines [GP0301]
Hydroxy fatty acids [FA0105]
Diacylglycerophosphoethanolamines [GP0201]
Unsaturated fatty acids [FA0103]
Dicarboxylic acids [FA0117]
Oxo fatty acids [FA0106]
Diacylglycerophosphates [GP1001]
phytoceramides [SP0203]
number of peaks dysregulated in METTL7B-KO cells (%)

**Figure S3 | METTL7B knockout reproducibly alters lipid composition across lipidomics and metabolomics in negative and positive ion modes.**

**a** Scatter plot showing the correlation of $\log_2$ fold changes (KO/WT) for each detected peak between two independent METTL7B-KO clones in four datasets: lipidomics (negative ion mode), lipidomics (positive ion mode), metabolomics (negative ion mode), and metabolomics (positive ion mode). Each point represents a peak; colors indicate point density. Spearman's correlation test. **b** Alluvial plots showing the lipid class composition of peaks in each dataset (lipidomics/metabolomics, negative/positive ion modes). For each mode, the proportions of lipid classes are shown separately for upregulated peaks (up), all detected peaks (all), and downregulated peaks (down) in METTL7B-KO cells. **c** Stacked bar charts showing the percentage of dysregulated peaks within each main lipid class for each detection mode (lipidomics/metabolomics, negative/positive ion modes). Each bar is segmented into downregulated (blue), unchanged (gray), and upregulated (red) portions. **d** Stacked bar charts showing the percentage of dysregulated peaks within each lipid subclass for each detection mode. Each bar is segmented as in panel C.

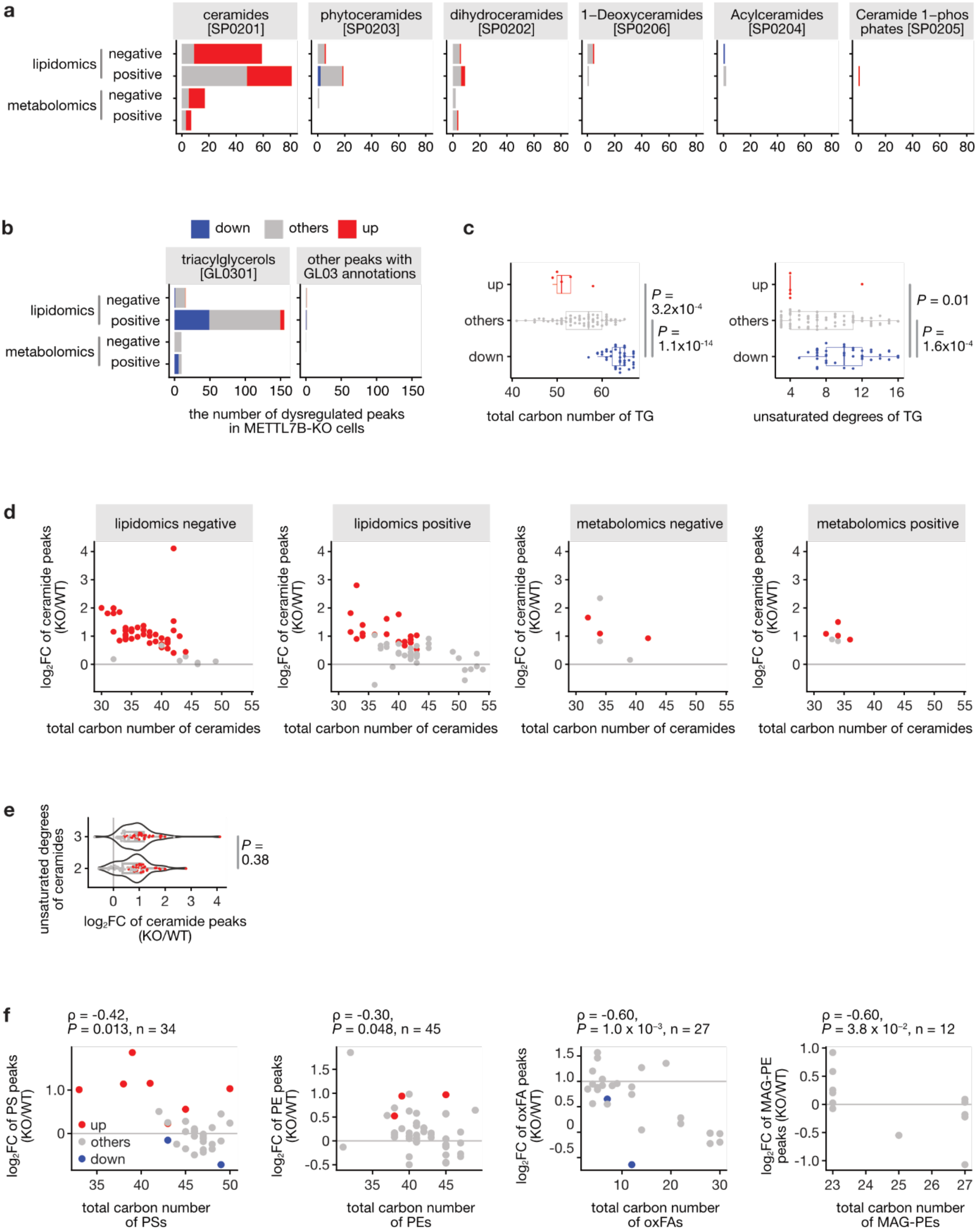
a
ceramides [SP0201]
phytoceramides [SP0203]
dihydroceramides [SP0202]
1−Deoxyceramides [SP0206]
Acylceramides [SP0204]
Ceramide 1−phos phates [SP0205]
lipidomics
metabolomics
negative
positive
b
down
others
up
triacylglycerols [GL0301]
other peaks with GL03 annotations
the number of dysregulated peaks in METTL7B-KO cells
c
total carbon number of TG
unsaturated degrees of TG
P = 3.2x10⁻⁴
P = 1.1x10⁻¹⁴
P = 0.01
P = 1.6x10⁻⁴
d
lipidomics negative
lipidomics positive
metabolomics negative
metabolomics positive
log₂FC of ceramide peaks (KO/WT)
total carbon number of ceramides
e
unsaturated degrees of ceramides
log₂FC of ceramide peaks (KO/WT)
P = 0.38
f
ρ = -0.42, P = 0.013, n = 34
ρ = -0.30, P = 0.048, n = 45
ρ = -0.60, P = 1.0 x 10⁻³, n = 27
ρ = -0.60, P = 3.8 x 10⁻², n = 12
log₂FC of PS peaks (KO/WT)
log₂FC of PE peaks (KO/WT)
log₂FC of oxFA peaks (KO/WT)
log₂FC of MAG-PE peaks (KO/WT)
total carbon number of PSs
total carbon number of PEs
total carbon number of oxFAs
total carbon number of MAG-PEs

**Figure S4 | METTL7B knockout affects triacylglycerol and ceramide species across detection modes.**

**a** Stacked barplots showing the number of detected peaks for triacylglycerols [GL0301] (left) and other peaks with triradylglycerol [GL03] annotations (right) across four detection modes (lipidomics/metabolomics, negative/positive ion modes). Each bar is segmented into downregulated (blue), unchanged/other (gray), and upregulated (red) peaks. **b** Stacked barplots showing the number of detected peaks for ceramide subclasses—ceramides [SP0201], phytoceramides [SP0203], dihydroceramides [SP0202], 1-deoxyceramides [SP0206], acylceramides [SP0204], and ceramide 1-phosphate [SP0205]—across four detection modes. Each bar is segmented as in panel a. **c** Box plots showing the distribution of total carbon numbers (left) and degrees of unsaturation (right) of triacylglycerol (TG) peaks classified as upregulated (red), others (grey), and downregulated (blue) peaks. Wilcoxon test. **d** Scatter plots showing $log_2$FC (KO/WT) of ceramide peaks versus their total carbon number, separated by detection mode (lipidomics negative, lipidomics positive, metabolomics negative, metabolomics positive). Points are colored by regulation status (red: upregulated, blue: downregulated, grey: others). **e** Violin plots showing the distribution of $log_2$ fold change (KO/WT) of ceramide peaks grouped by degree of unsaturation. Wilcoxon test. **f** Scatter plots showing $log_2$FC (KO/WT) of phosphatidylserine (PS) peaks, phosphatidylethanolamine (PE) peaks, oxidized fatty acids (oxFA) peaks, and monoacylglycerophosphoethanolamine (MAG-PE) peaks versus their total carbon number. Points are colored by regulation status (red: upregulated, blue: downregulated, grey: others). Spearman's correlation test.
